# Effects of lowered [Na^+^]_o_ and membrane depolarization on the Ca^2+^ transients of fast skeletal muscle fibers. Implications for muscle fatigue

**DOI:** 10.1101/2022.04.28.489961

**Authors:** Marbella Quiñonez, Marino DiFranco

## Abstract

Sodium (Na^+^) and potassium (K^+^) movements during repetitive stimulation of skeletal muscle fibers leads to lowered transmembrane Na^+^ and K^+^ gradients. Impaired calcium release resulting from the predicted reduction of the action potential (AP) overshoot (OS) has been suggested as a causative factor of muscle fatigue.

To test this hypothesis, we used a double grease-gap method and simultaneously recorded membrane action potentials (MAPs) and Ca^2+^ release (as Ca^2+^ transients), elicited by single pulses or short trains of pulses (100 Hz, 100 ms), in rested fibers polarized to membrane potentials (Vm) between -100 to -55 mV, and exposed to various extracellular Na^+^ concentrations ([Na^+^]_o_; 115, 90, 60 and 40 mM).

In single stimulation experiments, we found that at physiological Vm (-100 mV), Ca^2+^ release was mostly immune to [Na^+^]_o_ reductions up to 60 mM (~1/2 the physiological value). In contrast, at 40 mM Na^+^_o_ Ca^2+^ release was reduced by 80%, notwithstanding robust MAPs with large OS (~30 mV) were recruited in this conditions.

At Vm between -100 and -60 mV, a 20% reduction of [Na^+^]_o_ (115 to 90 mM) had no major detrimental effects on Ca^2+^ release. Instead, depolarization-dependent potentiation of Ca^2+^ transients, with a maximum at -65 mV, was observed at both 115 and 90 mM Na^+^_o_. Potentiation was smaller at 90 mM Na^+^_o_. At both [Na^+^]_o_, maximally potentiated Ca^2+^ transients (i.e. at -60 mV) were recruited by MAPS with reduced OSs.

In contrast, Ca^2+^ release was significantly depressed and no potentiation was observed at Vm between -100 to -70 mV when [Na^+^]_o_ was reduced 60 mM.

At extreme Na^+^_o_ (40 mM), Ca^2+^ release recorded at Vm between -100 and -70 mV was almost obliterated; nonetheless robust MAPs, with OSs of ~25 mV, were recruited.

Extreme depolarizations significantly depressed Ca^2+^ release at all [Na^+^]_o_ tested. The Vm leading to Ca^2+^ release depression was more negative the lower the [Na^+^]_o_ (-55, -60 and -70 for 115, 90 and mM Na^+^_o_, respectively).

Fiber exposed to 115-60 mM Na^+^_o_ can sustain normal Ca^2+^ release at a frequency of 100 Hz when polarized between -100 and -80 mV. Depolarizations beyond -80 mV lead to impaired Ca^2+^ release along the trains. In most cases, there was no correlation between changes in Ca^2+^ release and changes in OS. At 40 mM Na^+^_o_, only the 1st-3rd stimuli of trains recruited Ca^2+^ transients, which were significantly depressed vis a vis close to normal MAPs.

Neither the OS nor the duration of MAPs are figures of merit predicting the amplitude of Ca^2+^ transients. At critical combinations of depolarization, [Na^+^]_o_, and stimulation frequency, potentiated Ca^2+^ transients are recruited by MAPS with small OSs; and conversely, partial or total decoupling of Ca^2+^ release from close to normal MAPs was observed.

Depolarization and Na^+^_o_ deprivation depressed Ca^2+^ release in a synergistic way; lowered [Na^+^]_o_ increased the detrimental effects of depolarization on Ca^2+^ release, and depolarization render the ECC process more sensitivity to Na^+^_o_ deprivation.

Impaired TTS AP generation and/or conduction may explain the detrimental effects of depolarization and Na^+^_o_ deprivation on Ca^2+^ release.

The effects of increased K^+^_o_ and Na^+^_o_ deprivation on the force generation of rested fibers can be explained on the basis of the effects of membrane depolarization and Na^+^_o_ deprivation on Ca^2+^ release.

**Definitions:** [ion]_i_, [ion]_o_: intracellular and extracellular ion concentrations; ion= Na^+^, K^+^, Ca^+2^. (in molar units)

EFM-Na, EMF-K: electromotive force of Na^+^ and K^+^ (in mV)

ENa, EK: equilibrium potential for Na^+^ and Na^+^ (in mV)

Vm: membrane or holding potential (in mV)

TTS: transverse tubular system.

Ca-FWHM, Ca^+2^ transient full-width at half-maximum (in ms)

MAP-FWHM: MAP full-width at half-maximum (in ms)

REF: releasing effective time, time a MAP waveform is above -40 mV (in ms)

## Introduction

Repetitive stimulation of skeletal muscle fibers leads to a gradual decline of their capacity to generate mechanical work, a phenomenon referred to as muscle fatigue [1], for a review, see [2]. Although the exact mechanisms underlying muscle fatigue are still debated, membrane depolarization, and changes in the extra- and intracellular K^+^ and Na^+^ concentrations (**[K^+^]_o_, [Na^+^]_o_**, **[K^+^]_i_**, **[Na^+^]_i_**) resulting from repetitive stimulation has been long suggested as possible causatives factors [2, 3].

Impaired generation, waveform, or conduction of Na-dependent action potentials (APs), the physiological trigger of the excitation-contraction coupling (**ECC**) process, have been invoked as mechanisms by which depolarization and changes in [K^+^]_o_ and [Na^+^]_o_ leads to fatigue.

The effects of increases in K^+^_o_ and reductions in K^+^_i_ on force generation have been proposed to be mediated by membrane depolarization ensuing from the resulting reduction in K^+^ electromotive force (**K-EMF**); which in turn will reduce the sodium conductance (**gNa**) and thus impair the generation and conduction of APs. The consequent depression of the ECC process would reduce the capacity to generate force [4, 5]. This explanation for the activity dependent impairment of force generation is usually referred to as the “potassium hypothesis” of muscle fatigue [4].

The detrimental effects of Na^+^_o_ deprivation and Na^+^_i_ accumulation on force generation are explained by a reduction in the Na^+^ electromotive force (**Na-EMF**) leading to a reduced AP overshoot (**OS**) and impaired AP conduction. This alterations will depress Ca^2+^ release and subsequent force generation. This explanation is referred to as the “sodium hypothesis” of muscle fatigue [6].

A practical experimental paradigm to assess the role of changes in [K^+^]_o_ or [Na^+^]_o_ on muscle fatigue generation has consisted in measuring the mechanical output of rested muscles or bundles of muscle fibers in response to field stimulation while the concentration of one ion is changed and the other is maintained constant [4, 6, 7]. It is assumed that factors contributing to fatigue development during repetitive stimulation, should also affect force development in rested fibers [4]. This experimental paradigm does not allow for amenable measurements of membrane potential, let alone its control. While this approach has been key to further our understanding of muscle fatigue, the number of studies assessing changes in intermediate steps of the ECC process (i.e. Ca^2+^ release) is scant.

We have extended this approach by using cut segments of single fibers under current clamp conditions and directly measuring Ca^2+^ release with an impermeant low affinity Ca^2+^-sensing dye. In these conditions, membrane potential and extracellular and myoplasmic ionic composition can be measured and/or controlled [7].

Using this method we previously tested directly some of the tenets of the potassium hypothesis [4]. In particular, we have demonstrated that a)changes in Ca^2+^ release in response to increased [K^+^]_o_ up to, but not beyond, 10 mM are mediated by membrane depolarization and raised resting intracellular calcium concentration (**[Ca^2+^]_i_**); b)the effects of [K^+^]_o_ changes on Ca^2+^ release can be mimicked or counteracted by membrane potential changes imposed by current injection; and, c)depolarization-dependent (and [K^+^]_o_-dependent) changes in action potential overshoot and duration cannot individually explain the changes in Ca^2+^ release in response to membrane depolarization. It was concluded that the effects of elevated K^+^_o_ (and thus, depolarization) on the twitch force of rested muscles are mediated by its effects on Ca^2+^ release and that the concomitant changes in the AP waveform (i.e. OS, duration) do not directly correlate with changes in Ca^2+^ release [7].

The goal of the present work was to test the Na^+^ hypothesis for muscle fatigue. To this end, we studied the effects of Na^+^_o_ deprivation on the Ca^2+^ release of muscle fibers maintained at various membrane potentials and stimulated with single pulses or brief trains of pulses.

## Methods

### Animal model

Animals were handled in accordance to the regulations laid down by Universidad Central de Venezuela. Animals were euthanized by rapid transection of the cervical spinal cord followed by pithing in the cranial and caudal directions. Experiments were performed with segments of fibers dissected from the dorsal head of the semitendinosus muscle of tropical toads (Leptodactilus sp.).

### Electrophysiological techniques

Segments of fibers, cut at both ends, were mounted in an inverted double grease-seal chamber originally designed in our laboratory [7, 8] and maintained in current clamp conditions as previously described. The grease seals delimit three electrically isolated sections. The electrical activity and Ca^2+^ release are measured in the central section. The two lateral sections afford diffusional access to- and electrical contact with the myoplasm at the central section. The current- and voltage-clamp amplifier was home made. Experiments were started 20 min after fibers were mounted in the experimental chamber. Contraction was prevented by stretching the fibers to sarcomere lengths of about 4μm. This condition also allowed for the Ca^2+^ dye calibration at the end of experiments [7].

The mean diameter and sarcomere length were 62.8 ± 8 μm and 4.2 ± 0.5 μm, respectively (36 fibers, 9 toads).

The membrane potential (or holding potential, Vm) was initially adjusted to −100 mV, and then varied to values between –100 to –55 mV by manually adjusting the holding current.

Action potentials were elicited by single pulses or short trains of pulses (100 Hz, 10 pulses). Pulse duration was 0.2 ms, and amplitude was adjusted to ~15% above the threshold (determined from −100 mV), and not varied thereafter. Since regenerative responses are recruited simultaneously at all points of the segment of fiber in the central pool of the experimental chamber [7], non-propagated or membrane action potentials (MAPs) are elicited.

The fibers were continuously perfused with the desired solution. A 3 min period was allowed after varying the holding potential, and a 3 min period was allowed for equilibration after changing solutions. When repetitive stimulation was used, 2 min were allowed between consecutive trains. The use of large equilibration aim at imposing a desired [Na^+^] in the lumen of the t-tubules without radial gradients.

We measured the overshoot (OS) and duration of MAPs recorded at the different conditions used. The OS is, by definition, the difference between the peak of a MAP and 0 mV. The duration of MAP waveforms was measured as a)the full-width at half-maximum (**FWHM**) and b)the full-width at −40 mV. The later represents the time the membrane potential is above −40 mV, a typical value for the Ca^2+^ release threshold in voltage clamp conditions [9, 10]. This parameter represents, in practice, the time during which Ca^2+^ release can occur, i.e. the <u>r</u>eleasing <u>e</u>ffective <u>t</u>ime (**REF**). REF definition stems from the classical mechanically effective time [11, 12].

All experiments were performed at room temperature (21-22°C).

### Solutions

The central section of the fibers were bathed in Ringer solution (in mM: 115 NaCl, 2.5 KCl, 1.8 CaCl_2_, 1 MgCl_2_ and 10 4-morpholineethanesulfonic acid [MOPS]; titrated with NaOH) while both cut ends of the fibers were bathed in an “internal” solution designed to approximate the myoplasm ionic composition (in mM: 110 aspartate, 5 K_2_-ATP, 5 Na_2_-creatine phosphate, 20 MOPS, 0.1 ethylene glycol tetraacetic acid [EGTA], 5 MgCl_2_, titrated with KOH and added with 0.5 mg/ml creatine phosphokinase) [7]. Reductions in [Na^+^]_o_ were compensated by additions of N-methyl-D-glucamine, thus osmolarity was maintained constant. [Na^+^]_o_ was changed from 115 mM to 90, 60 and 40 mM. [K^+^]_o_ was maintained constant in all experiments (i.e. 2.5 mM). All solutions, had a pH=7.2 and an osmolarity of 250 ± 3 mOsmol/kg H_2_O. Assuming an intracellular Na^+^ activity of 10 mM (i.e. identical to that of the internal solution [7]), a temperature of 20°C (295°K), and ideal selectivity of the sodium channels, the equilibrium potential for Na^+^ (**ENa**) in the presence of 115, 90, 60 and 40 mM extracellular Na^+^ was calculated as 62, 56, 47 and 35 mV, respectively. All chemicals were from Sigma.

### Calcium measurement

Changes in [Ca^2+^]_i_ elicited by MAPs (thereafter referred as Ca^2+^ transients) were followed with a low affinity impermeant fluorescent calcium dye (Oregon green 488 BAPTA 5N, thereafter, OGB5N; ThermoFisher Scientific). Ca^2+^ dependent fluorescence transients were measured using the setup previously described [7]. [Ca^2+^]_i_ changes (i.e. in actual μM units) were calculated using the same parameters and methods described previously [7]. The peak (μM) and duration of Ca^2+^ transients recorded at different conditions were measured. Duration was evaluated as the full-width at half-maximum (**FWHM**), and referred to as **Ca-FWHM** thereafter.

### Data acquisition and analysis

Membrane potential and fluorescence signals were filtered at 5 and 2 kHz, respectively, using 8-pole Bessel filters (Frequency Devices); and acquired simultaneously using a Digidata 1200A acquisition system and Axotape software (Molecular Devices). Data were analyzed using Origin 8.0 (Origin Microcal). Data are presented as means ± SE. Means were compared using the Student’s t-test. Statistical significance was set at p<0.05.

## Results

### Effects of reducing [Na^+^]_o_ on Ca^2+^ transients recorded from fibers maintained at −100mV

We first studied the Ca^2+^ transients and MAPs elicited by single pulse stimulation in fibers maintained at a holding potential of −100 mV and chronically exposed to 115, 90, 60 and 40 mM Na^+^_o_ (Figure 1). In this way, the effects of changes in [Na^+^]_o_ on MAP generation and Ca^2+^ release can be studied at constant Vm and gNa. Also, at this potential the availability of Na^+^ channels and the voltage sensor for ECC is maximized.

**Figure 1.**
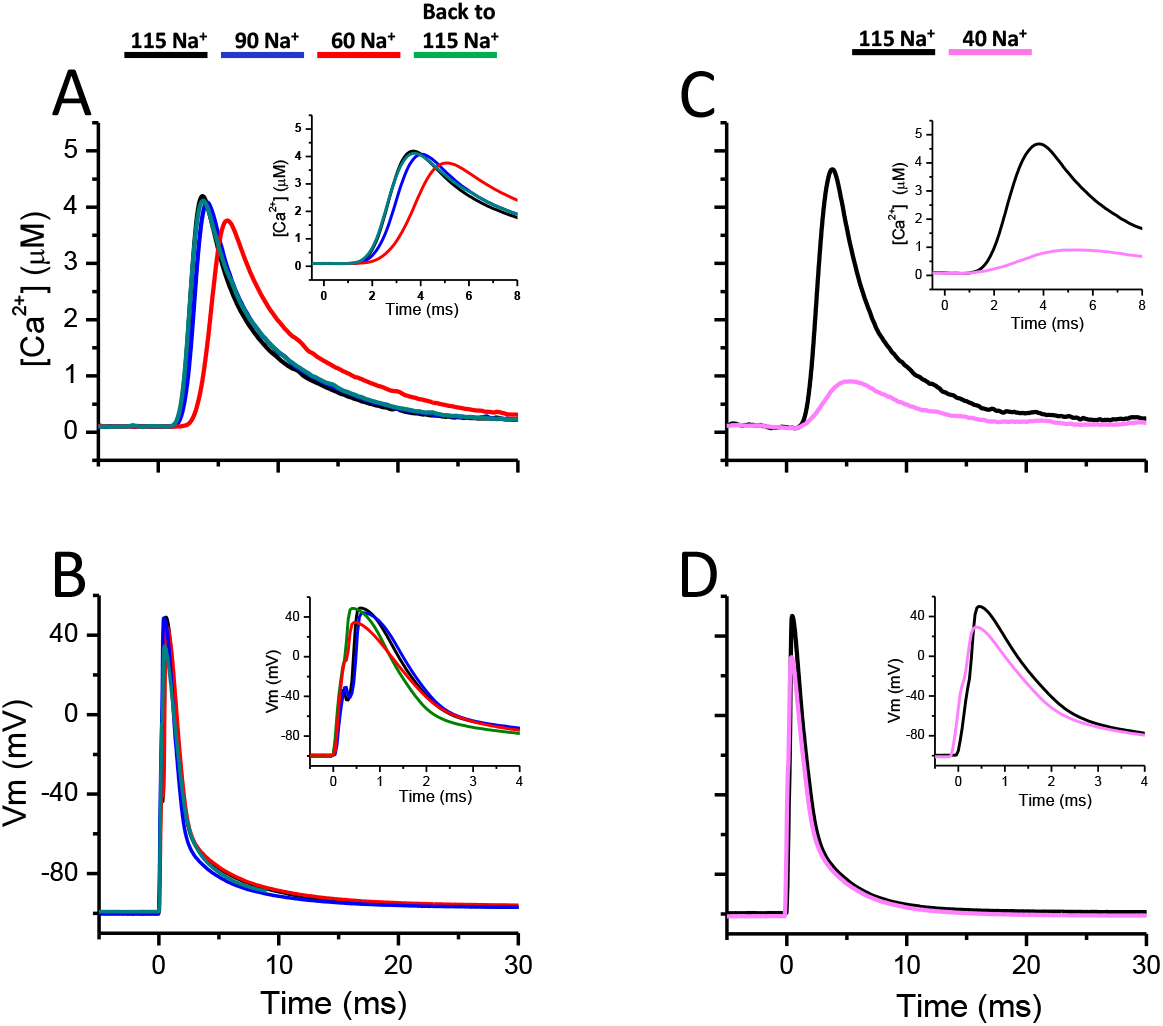
Ca^+2^ release elicited by single pulse stimulation in fibers maintained at − 100 mV and exposed to various [Na^+^]_o_. **A.** The black, blue and red traces represent, respectively, the Ca^+2^ transients recorded from a fiber exposed successively to at 115, 90 and 60 mM Na^+^_o_. The green trace represents the Ca^+2^ transient recorded after returning to 115 from 60 mM Na^+^_o_. **B.** MAPs recorded simultaneously with each of the Ca^+2^ transients shown in A, represented using the same color coding as in A. **C.** Ca^+2^ transients recoded in another fiber exposed successively to 115 (black trace) and 40 mM Na^+^_o_ (magenta trace). **D.** MAPs eliciting the Ca^+2^ transients shown in C. The same color coding as a in C was used. The inset in each panel shows the corresponding Ca^+2^ transients or MAPs in an expanded time scale. Notice that, for all panels, two different time scales are used in the insets for Ca^+2^ transients and MAPs.

A substantial 25 mM reduction of [Na^+^]_o_ from 115 to 90 mM results only in a slight reduction in the raising phase and a slight increase in the time to peak of Ca^2+^ transients (Figure 1A and inset, black and blue traces, respectively). Likewise, the simultaneously recorded MAPs eliciting those Ca^2+^ transients are very similar to each other (Figure 1B and inset, black and blue traces).

A further reduction of [Na^+^]_o_ to 60 mM (e.g. a 55 mM change) leads per se to a slightly smaller Ca^2+^ transient, with a delayed onset and a the longer time to peak as compared with records obtained at physiological [Na^+^]_o_ (Figure 1A and inset, red trace). These changes are accompanied by a small reduction in the MAP OS (Figure 1B). Changes induced by reducing Na^+^_o_ were fully reversed by returning to Ringer solution containing 115 mM Na^+^ (Figures 1A and 1B and insets, green traces). Reversibility was demonstrated for all other conditions used in this work (not shown).

The most dramatic effects are seen upon reducing [Na^+^]_o_ to 40 mM (close to 1/3 of the physiological value). As shown in Figure 1C and inset (magenta trace), the amplitude of the Ca^2+^ transient was reduced by about 80% in the presence of 40 mM [Na^+^]_o_, and the time to peak significantly increased. Notably, the changes in the corresponding MAP are relatively minor as compared with the gross impairment of the Ca^+^ transient. Although the MAP OS was reduced from 51 to 30 mV (Figure 1D and inset, cyan trace), this change seems in itself insufficient to explain the almost obliteration of the Ca^2+^ release (Figure 1D and inset, cyan trace). Since Vm was kept constant, gNa should have also remained constant; consequently, variations in the OS were small; as expected from the calculated ENa.

In contrast to what was observed in response to changes in [K^+^]_o_ [7], no significant changes in pre-stimulus [Ca^+^]_i_ were detected in response to reductions of [Na^+^]_o_ alone.

### Effects of membrane depolarization on Ca^2+^ transients recorded from fibers exposed to various [Na^+^]_o_

The previous section demonstrated that, in fibers maintained at −100 mV and exposed to 2.5 mM K^+^_o_, the Ca^2+^ release is practically immune to [Na^+^]_o_ reductions down to 60 mM. We have previously demonstrated that the effects of K-dependent depolarization on Ca^2+^ release can be mimicked by current injection [7]. In order to assess the combined effects of changes in resting membrane potential and [Na^+^]_o_ on the ECC process, we next recorded Ca^2+^ transients and MAPs elicited by single stimulation in fibers exposed to various [Na^+^]_o_ (115, 90, 60 and 40 mM) and maintained at potentials between −100 and −55 mV (typically, −100, −90, −80, −70, −65, −60 and −55 mV) by steady current injection while [K^+^]_o_ was maintained constant. Representative records obtained at each condition are shown in Figure 2.

**Figure 2.**
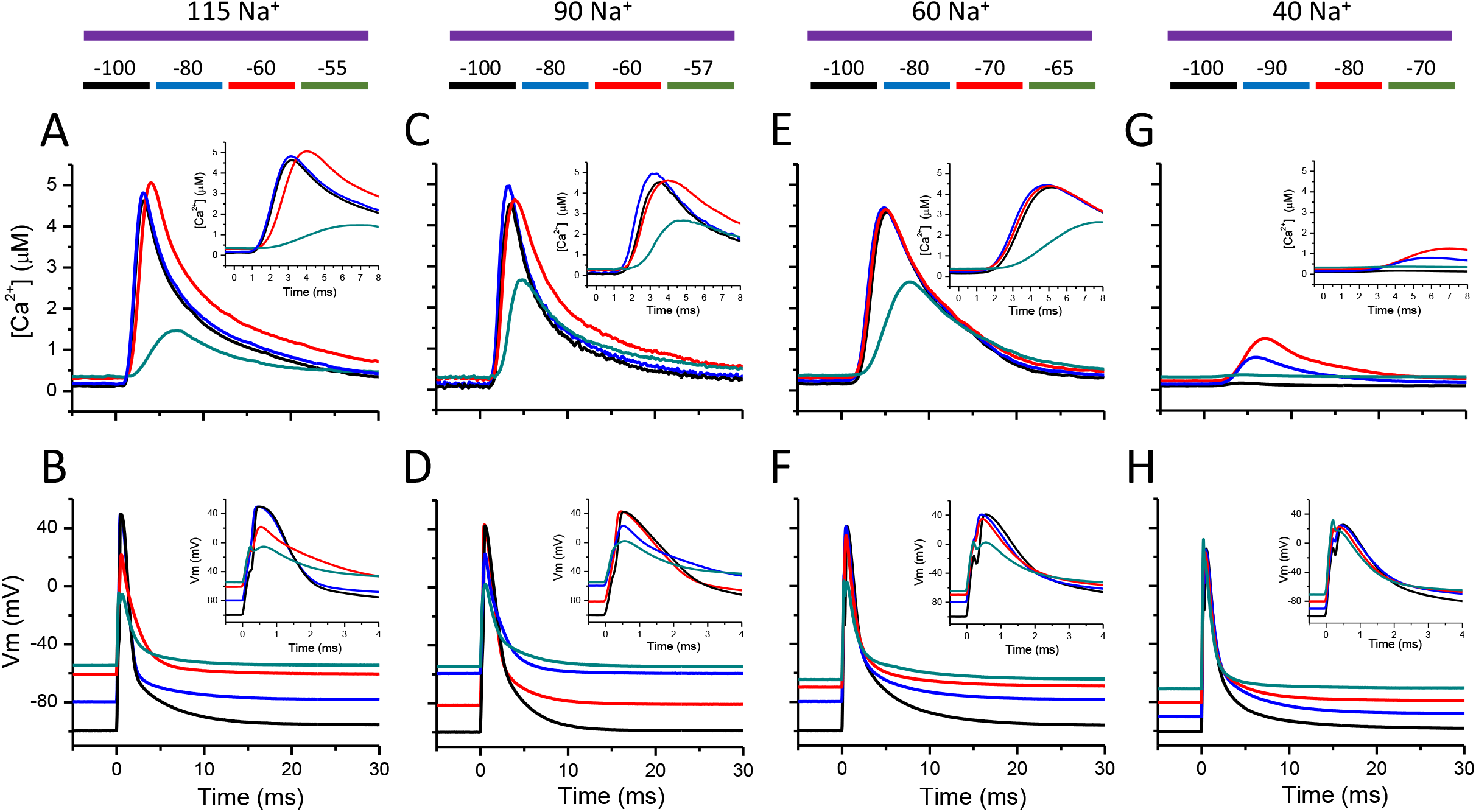
Effects of steady depolarization on the Ca^+2^ transients recorded from fibers exposed to various [Na^+^]_o_. Ca^+2^ transients (top panels) and their corresponding MAPs (bottom panels) recorded from fibers bathed in 115, 90, 60 and 40 mM Na^+^_o_, and maintained at various membrane potentials, as indicated by the color bars. **A-B**. Records at 115 mM Na^+^_o_. **C-D**. Records at 90 mM Na^+^_o_. **E-F**. Records at 60 mM Na^+^_o_. **G-H**. Records at 40 mM Na^+^_o_. For each [Na^+^]_o_, different holding potentials are used, records at each condition are color coded. The same color is used for simultaneously recorded Ca^+2^ transients and MAPs. The inset in each panel shows the corresponding Ca^+2^ transients or MAPs in an expanded time scale. Notice that, for all panels, two different time scales are used in the insets for Ca^+2^ transients and MAPs.

As previously reported [7], in the presence of 115 mM Na^+^_o_, imposed steady membrane depolarizations has a dual effect on Ca^2+^ release. Ca^2+^ transients are potentiated for depolarizations between −100 and −60 mV (Figure 2A and inset, black, blue and red traces), but their amplitude is sharply reduced for further depolarizations to −55 mV (Figure 2A and inset, green trace). Fiber depolarization led to an increase in resting [Ca^2+^]_i_, as previously shown [7].

MAPs eliciting the Ca^2+^ transients in Figure 2A are shown in Figure 2B. MAPs recorded at −100 and −80 mV differ only in the imposed Vm (Figure 2B and inset, black and blue traces). Instead, significant reductions in the OS are seen for depolarizations to −60 and −55 mV. Notably, a mere 5 mV depolarization leads from the more potentiated Ca^2+^ transient (at −60 mV, Figure 2B and inset, red trace) to the more depressed one (Figure 2B and inset, green trace). These changes may probably results from voltage dependent reduction of gNa.

Similar responses were obtained for fibers exposed to 90 mM Na^+^_o_ (Figure 2C and 2D). The extreme sensitivity of Ca^2+^ release to membrane potential is exemplified by records obtained at −57 mV (Figure 2C and 2D, green traces). Both the values of peak Ca^2+^ release and the OS are larger than those obtained at −55 mV in the presence of 115 mM Na^+^_o_.

Ca^2+^ release is well maintained in the presence of 60 mM Na^+^_o_ over a range of membrane potentials spanning from −100 to −70 mV. Nonetheless, in these conditions, aside from a reduced amplitude of the Ca^2+^ transients, interesting changes in the ECC are observed. The depolarization-dependent potentiation of Ca^2+^ release seen in fibers exposed to 115 and 90 mM Na^+^_o_ is absent in the presence of 60 mM Na^+^_o_, i.e. Ca^2+^ release is essentially identical at membrane potentials between −100 and −70 mV (Figure 2E and inset, black, blue and red traces). In addition, depolarization-dependent depression of ECC is shifted leftward, as a significant reduction in peak Ca^2+^ release is seen at −65 mV (Figure 1E and inset, green trace), a membrane potential that elicits potentiation in fibers bathed in 115 and 90 mM Na^+^_o_. In these last conditions, the AP essentially lacks an OS (Figure 2F and inset, green trace). These results clearly show that sensitivity to depolarizations is increased at reduced [Na^+^]_o_.

Ca^2+^ release was almost abolished at all membrane potentials tested when fibers are equilibrated in 40 mM Na^+^_o_ (Figure 2G and 2H). Interestingly, while Ca^2+^ release is barely distinguishable from baseline in fibers polarized to either −100 or −70 mV (Figure 2G and inset, black and green traces), sizable Ca^2+^ transients are detected at −90 and −80 mV (Figure 2G and inset, blue and red traces). Figure 2H shows that MAPs elicited from −100 and −70 mV display a large OS (about 25 mV) still failed to recruit sizable Ca^2+^ release (black and green traces). Also, MAP elicited from −90 and −80 mV have a similar OS and yet the Ca^2+^ release at −80 mV is larger than that at −90 mV (Figure 2H and inset, blue and red traces). These data show that at very low values, [Na^+^]_o_ has dominant depressing effects on the ECC process, suggesting a decoupling between mostly normal MAPS and Ca^2+^ release.

### Voltage dependence of Ca^2+^ release in fibers exposed to various [Na^+^]_o_

Experiments similar to those in Figure 2 were conducted in a population of fibers (see Figure 3) to assess how combined changes in [Na^+^]_o_ and membrane potential affect Ca^2+^ release. To get insight on the dependence of Ca^2+^ release on resting potential alone we first plotted the peak of Ca^2+^ transients recorded at each [Na^+^]_o_ as a function of Vm (Figure 3A).

**Figure 3.**
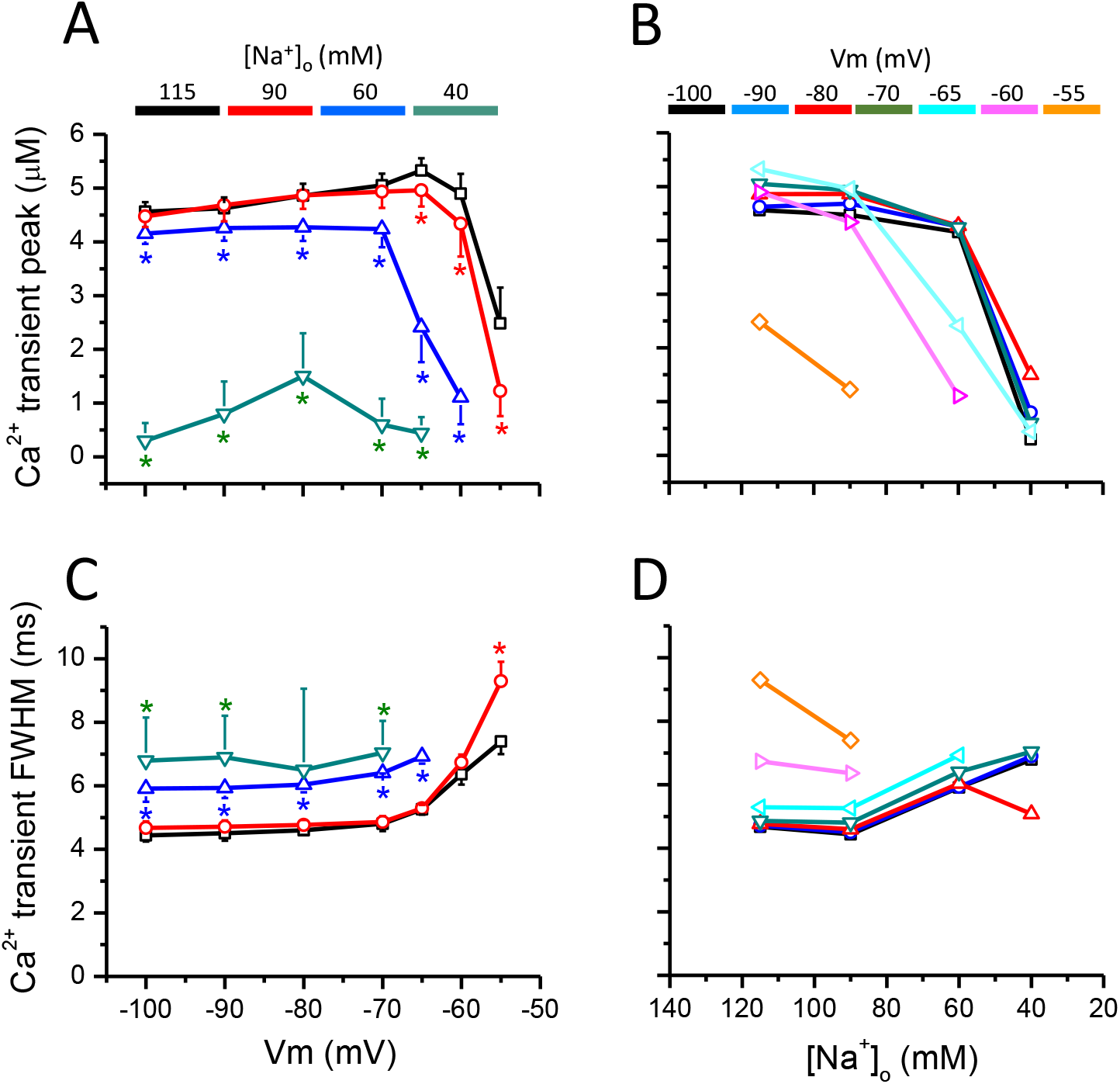
Dependence of Ca^+2^ transients’ peak and duration on the membrane potential and [Na^+^]_o_. **A.** Voltage dependence of the Ca^+2^ transient’s peak. Data points represent the means of Ca^+2^ transients’ peaks recorded at −100, −90, −80, −70, −60 and −55 mV from fibers exposed to 115 (black line and symbols), 90 (red line and symbols), 60 (blue line and symbols) and 40 mM Na^+^_o_ (green line and symbols). **B.** Dependence of the Ca^+2^ transient’s peak on [Na^+^]_o_. The data in A are plotted as a function of [Na^+^]_o_, the error bars are omitted for clarity. The holding potential (−100, −90, −80, −70, −65, −60 and −5 mV) is color coded as indicated by the colored horizontal bars. **C.** Voltage dependence of the Ca-FWHM. Data points represent the means of FWHM of Ca^+2^ transients recorded from fibers maintained at −100, −90, −80, −70, −60 and −55 mV and exposed to 115 (black line and symbols), 90 (red line and symbols), 60 (blue line and symbols) and 40 mM Na^+^_o_ (green line and symbols). Same population of fibers as in A. **D.** Dependence of the Ca-FWHM on the [Na^+^]_o_. The data in C are plotted as a function of [Na^+^]_o_, the error bars are omitted for clarity. The holding potential used is color coded as indicated by the color horizontal bars. Symbols represents mean ± SE. n=7, 6, 5, 3 for experiments at 115, 90, 60 and 40 mM Na^+^_o_. Asterisks indicate statistical difference from data at −100 mV.

The first significant finding is the resilience of Ca^2+^ release to large depolarizations even in face of [Na^+^]_o_ reductions down to 60 mM (Figure 3A, black, red and blue traces). In fact, fiber depolarizations from −100 mV similarly potentiate Ca^2+^ release for both 115 and 90 mM Na^+^_o_, with a maximum effect at −65 mV (Figure 3A, black and red traces). Potentiation is not seen in fibers bathed in 60 mM [Na^+^]_o_; instead a relatively small, almost voltage-independent reduction in the peak of Ca^2+^ transients was observed for depolarizations up to −70 mV, as compared with those recorded at 115 mM Na^+^_o_ (Figure 1A, black and blue traces).

The second finding is that Ca^2+^ release is significantly reduced when fibers are depolarized beyond −70 or −60 mV, depending on the [Na^+^]_o_. At high [Na^+^]_o_ (i.e. 115 and 90 mM, black and red traces, Figure 3A), Ca^2+^ release is reduced to a large extent only for very large depolarizations to −55mV, while a large depression is seen for smaller depolarizations (i.e. beyond −70 mV, blue trace, Figure 3A) when fibers are bathed in 60 mM Na^+^_o_. Clearly, Ca^2+^ release is more sensitive to depolarization at lower [Na^+^]_o_.

A rather intriguing voltage-dependence of Ca^2+^ release is seen at extremely low [Na^+^]_o_. At 40 mM Na^+^_o_, Ca^2+^ release is largely reduced at all membrane potentials tested, with an apparent minimum reduction at −80 mV.

To better appreciate the dependence of Ca^2+^ release on [Na^+^]_o_, we plotted the peak of Ca^2+^ transients shown in Figure 3A as a function of [Na^+^]_o_ (Figure 3B).

This representation clearly shows that Ca^2+^ release elicited by single stimulation is very insensitive to reductions of [Na^+^]_o_ to about half the physiological concentration (i.e. 115 to 60 mM) as long as membrane potential is maintained between −100 and −70 mV (Figure 3B, black, blue, red and green traces). Nonetheless, for this membrane potential range, Ca^2+^ release is largely reduced if [Na^+^]_o_ is further decreased to 40 mM (Figure 3B, black, blue, red and green traces).

The sensitivity of Ca^2+^ release to reductions in [Na^+^]_o_ highly increases when fibers are further depolarized beyond −70 mV. At resting potentials of −65 to −60 mV, reducing [Na^+^]_o_ below 90 mM results in reduced Ca^2+^ release (Figure 3B, cyan and magenta traces), while at −55 mV further impairing of Ca^2+^ release is seen at 90 mM Na^+^_o_ (Figure 3B, orange trace). This data show that the larger the depolarization, the larger the effects of Na^+^_o_ deprivation.

For the same population of fibers, we also looked at the effects of depolarization and [Na^+^]_o_ deprivation on the duration (FWHM) of Ca^2+^ transients (thereafter, **Ca-FWHM**; Figures 3C and 3D).

In the range of membrane potentials between −100 and −70 mV, the duration of Ca^2+^ transients recorded from fibers bathed in 115 and 90 mM Na^+^_o_ were similar to each other (~4.5 ms) and almost insensitive to depolarization (Figure 3C, black and red traces). For further depolarizations to −55 mV, the FWHM of Ca^2+^ transients at 115 and 90 mM Na^+^_o_ increased significantly to about 7.7 and 9.5 ms, respectively (Figure 3C, black and red traces).

The Ca-FWHM recorded at 60 mM Na^+^_o_ were larger than those at 115 mM Na^+^_o_ at all membrane potentials in the range of −100 to −65 mV. In these conditions, the Ca-FWHM was mildly sensitive to depolarization (Figure 3C, blue trace).

The Ca^2+^ transients of fibers bathed in 40 mM Na^+^_o_ are significantly prolonged (~7 ms) as compared with those recorded at 115 mM Na^+^_o_, and mostly insensitive to membrane depolarization between −100 and −70 mV Figure 3B, green trace).

The dependence of Ca-FWHM on Na^+^_o_ is shown in Figure 3D. Reducing Na^+^_o_ from 115 to 90 mM had no significant effects on Ca^2+^ transients recorded in fibers polarized between −100 and −65 mV (Figure 3D, black, blue, red, green and cyan traces). For those same potentials, the Ca^2+^ transients were prolonged when [Na^+^]_o_ was further reduced to 40 mM. At 115 and 90 mM Na^+^_o_, highly depolarized fibers (−60 and −55 mV) showed the longest Ca^2+^ transients (Figure 3D, magenta and orange traces).

### Effects of membrane depolarization and reduction of [Na^+^]_o_ on the overshoot and FWHM of MAPs

In physiological conditions the ECC process is triggered by a longitudinally and radially propagated AP. It can be expected that either the amplitude or the width of the AP, or both parameters, may be determinant factors of the features of Ca^2+^ transients, or hence those of the twitch force. In contrast to physiological conditions, in our case, MAPs are the trigger of the ECC progress.

We first determined the dependence of the OS on membrane potential and [Na^+^]_o_. MAPs elicited by single stimulation in rested fibers polarized at −100 mV and exposed to 115 mM Na^+^_o_ have a mean OS close to 50 mV (the predicted ENa is 62 mV). Fiber depolarization and Na^+^_o_ deprivation are expected reduce the OS by different mechanisms. To study the extent of these effects, we plotted the OS as a function of the resting potential (Figure 4A) or [Na^+^]_o_ (Figure 4B).

**Figure 4.**
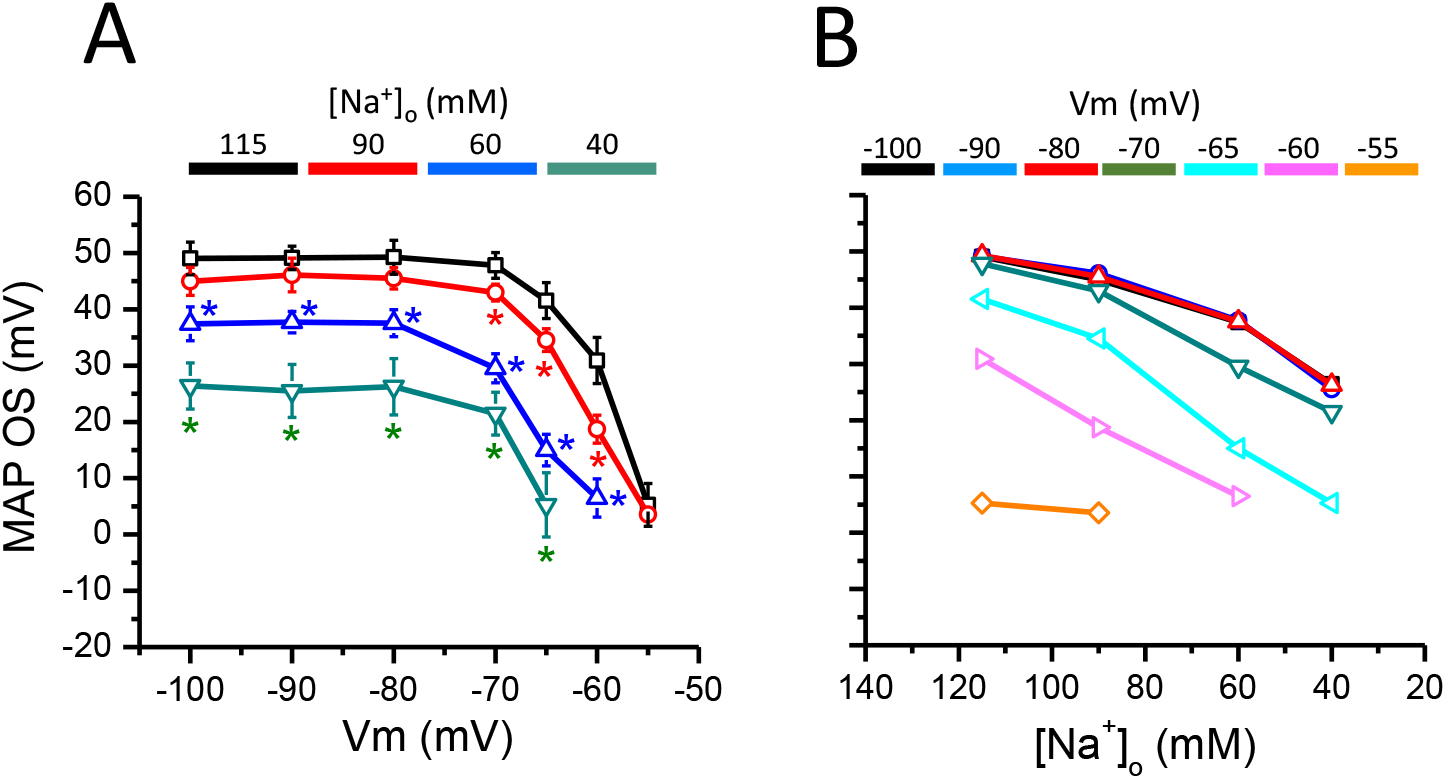
Dependence of MAP overshoot on holding potential and [Na^+^]_o_. **A.** The OS is plotted as a function of the membrane potential. Data points are the mean OS determined from MAPs recorded at −100, −90, −80, −70, −60 and −55 mV from fibers bathed in 115 (black line and symbols), 90 (red line and symbols), 60 (blue line and symbols) and 40 mM N (green line and symbols). **B.** Dependence of the OS on the [Na^+^]_o_. Data in A are plotted as a function of [Na^+^]_o_. Error bars are omitted for clarity. Each plot represent data obtained at the same holding potential (−100, − 90, −80, −70, −60 and −55 mV), as indicated by the colored horizontal bars. Symbols are mean ± SE. Asterisks indicate statistical difference from data at −100 mV.

Each plot in Figure 4A represents data obtained at a particular Na^+^_o_. For 115 and 90 mM [Na^+^]_o_, the OS was mostly insensitive to depolarizations along a 30 mV range from −100 to −70 mV, but drop abruptly to values close to zero when fibers are further depolarized to −55 mV, i.e. an additional 15 mV change (Figure 4A, black and red traces and symbols). Notably, in the range of potentials leading to Ca^2+^ release potentiation, the OS was either constant or decreased (compare Figures 3A and 4A). A similar voltage dependence was seen for 60 and 40 mM Na^+^_o_, but the transition point from voltage-independent to voltage-dependent OS values seems to be −80 mV, instead of −70 mV, as seen at larger [Na^+^]_o_ (Figure 4A, blue and green traces and symbols). Comparisons among the four plots in Figure 4A show that for any given membrane potential the lower the [Na^+^]_o_, the smaller the OS. The dependence of the OS on [Na^+^]_o_ is better demonstrated in Figure 4B; in this case, each plot represents data obtained at the same holding potential. For all membrane potentials explored, the OS decreases as the Na^+^_o_ is lowered from 115 to 40 mM. The dependence is close to linear if [Na^+^]_o_ is represented in a logarithmic scale (not shown). Noticeable, lowering [Na^+^]_o_ produced similar reductions on the OS at membrane potentials −100 and −80 mV, as demonstrated by the close overlapping of the black, blue and red traces in Figure 4B. However, the same changes in [Na^+^]_o_ produced much larger reductions in the OS for larger depolarizations to 60 mV (Figure 4B, cyan and magenta lines and symbols). At extreme depolarizations (i.e. −55 mV) the OS is little affected upon reducing [Na^+^]_o_ from 115 to 90 mM (Figure 4B, orange line and symbols).

We next measured the FWHM of MAPs (thereafter, **MAP-FWHM**) recorded from the same population of fibers (Figure 5A). It can be seen that for the four [Na^+^]_o_ tested, the MAP-FWHM decreases monotonically as fibers are depolarized from −100 to −70 mV. In addition, for 115 and 90 mM Na^+^_o_, further depolarization towards −55 mV resulted in a significant prolongation of MAPs. Moreover, for any given Vm between −100 and 70 mV, significantly wider MAPs are recorded at 90 and 60 Na^+^_o_ as compared with those at 115 Na^+^_o_. Overall, for 115 to 60 mM Na^+^_o_, voltage-dependence of changes in MAP-FWHM (Figure 5A) and Ca^2+^ release (Figure 3A) do no correlate.

**Figure 5.**
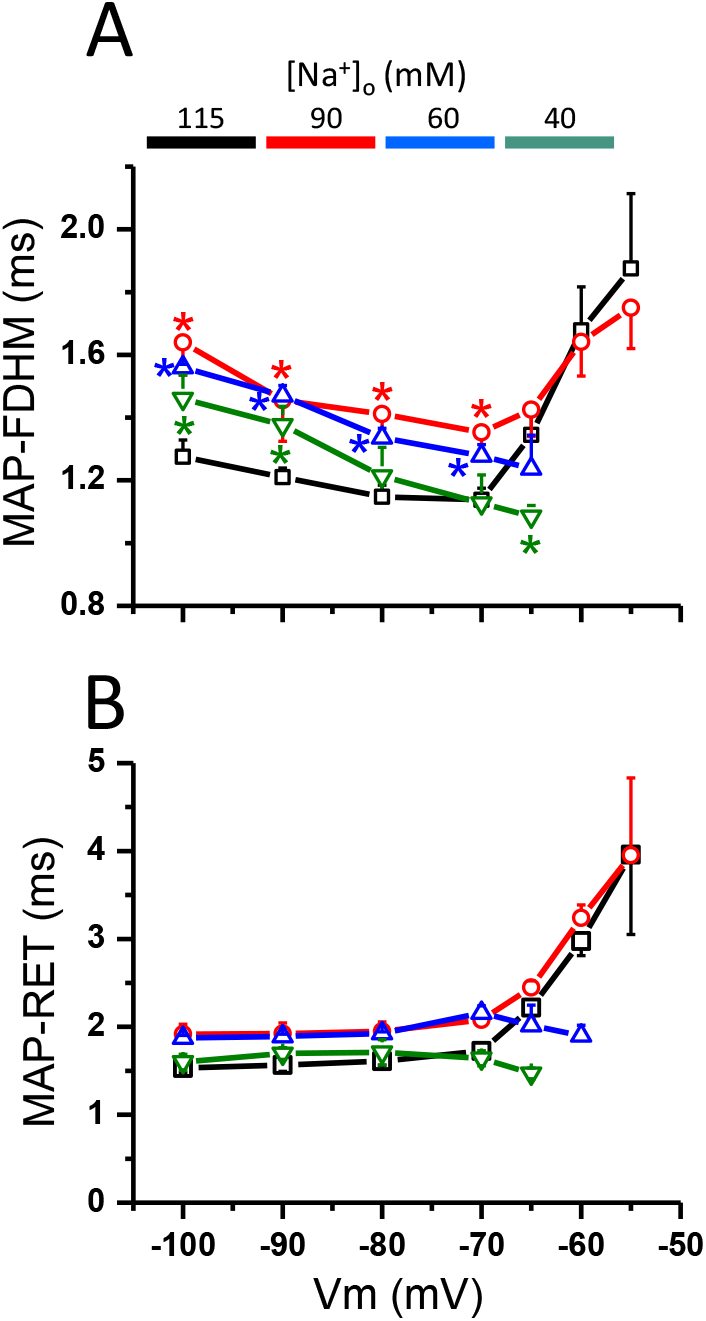
Voltage dependence of MAP FWHM and DUR-40. A. The FWHM of MAPs recorded in fibers exposed to various [Na^+^]_o_ is plotted as a function of holding potential (Vm). The [Na^+^]_o_ (115, 90, 60 and 40 mM) is color coded as indicated by the colored bar. B. The RET of MAPs recorded in fibers exposed to various [Na^+^]_o_ is plotted as a function of holding potential (Vm). The [Na^+^]_o_ (115, 90, 60 and 40 mM) is color coded as indicated by the colored bar. Symbols are mean ± SE. Asterisks indicate statistical difference from data at −100 mV.

We reasoned that this may be due to the fact that MAP-FWHM is measured at potentials more positive than the threshold for Ca^2+^ release in voltage clamp conditions, typically about −40 mV [9, 10]. Looking for a parameter that better correlates with the peak Ca^2+^ transients, we measured the RET of MAPs (thereafter, **MAP-RET**; Figure 5B, see also Methods for definition).

As expected, MAP-RET was larger than MAP-FWHM at all equivalent combinations of membrane potentials and [Na^+^]_o_. In contrast to MAP-FWHM, MAP-RET for all [Na^+^]_o_ is mostly independent of membrane depolarization in the range of −100 to −70 mV. For 90 and 60 mM Na^+^_o_, significantly larger MAP-RETs were measured from MAPs recorded at all membrane potential tested as compared with those measured at 115 mM Na^+^_o_. Thus, at least in the range of membrane potentials between −10 and −70 mV, MAP-RET better correlates with Ca^2+^ release than MAP-FWHM. At potentials positive to −70 mV both parameters negatively correlate with Ca^2+^ release.

### Effects of low [Na^+^]_o_ and depolarization on Ca^+2^ release elicited by repetitive stimulation

It is well stablished that tetanic force is more sensitive to [Na^+^]_o_ deprivation and depolarization than twitch tension [11], but similar studies of Ca^2+^ release are missing. Thus, we measured Ca^2+^ transients elicited by short (100ms) 100 Hz trains in rested fibers exposed to 40-125 mM Na^+^_o_ and maintained at holding potentials between −100 and −55 mV. Experiments in each [Na^+^]_o_ were repeated in 3-4 different fibers.

#### 115 mM Na^+^_o_

In physiological conditions (i.e. 115 mM Na^+^_o_ and −100 mV), when fibers were stimulated with short trains of pulses (100 ms) applied at 100 Hz, a distinct Ca^2+^ transient was recruited by each of 10 stimulus applied, i.e. fidelity equals 1 (Figure 5A, top panel). Nonetheless, regardless all the corresponding MAPs have an identical OS (~50 mV, Figure 5A, bottom panel), the peak of the first Ca^2+^ transient is not maintained along the trains. Instead, the peak of the Ca^2+^ transients decays from the first response, exceeding 5 μM, to a smaller, relatively stable value by the 4^th^ response, which is typically about 50% that of the first response. The interpulse [Ca^+2^]_i_ also remains elevated, at about 1.5 μM, along the trains as compared with the pre-stimulus value (Figure 4A, top panel). In contrast to the constancy of the OS, the interpulse membrane potential becomes progressively more positive along the train, reaching a relatively stable value by the end of stimulation.

Aside from the potentiation of the first Ca^2+^ transient (as seen in single stimulation experiments), at 115 Na^+^_o_, depolarization to −80 mV does not have significant effects on the Ca^2+^ release along the train (Figure 5B, top panel). The electrical responses at −80 mV are also comparable to those elicited at −100 mV (Figure 5B, bottom panel).

Depolarization to −60 mV also results in a predictable potentiation of the first Ca^+2^ transient, but an unexpected significant depression of the Ca^+2^ release was seen afterwards. While fidelity was no reduced, the Ca^+2^ transient’s peaks decay in an irregular fashion along the train, sometimes resembling alternants (Figure 5C, top panel). Interestingly, regardless the corresponding first MAP at −60 mV is reduced as compared to those at −100 and −80 mV, it still recruited a potentiated Ca^+2^ transient. While the rest of MAPs are smaller, they show similar features among them (Figure 5C, bottom panel); this do not correlates with the irregular changes observed in Ca^+2^ release peaks (Figure 5C, bottom panel).

Concurring with data from single stimulation, Ca^+2^ release is almost abolished at −55 mV (Figure 5D, top panel). In contrast, regular MAPs, with similar relatively large OSs of about 5 mV, were elicited by each pulse along the train (Figure 5D, bottom panel). Only a small Ca^+2^ transient was elicited by the first MAP, while the rest of MAPs failed to recruit any Ca^+2^ release.

Similar responses were confirmed in another 4 fibers.

#### 90-mM Na^+^_o_

Representative responses of fibers (n=4) bathed in 90 mM Na^+^_o_ to repetitive stimulation are shown in Figure 5E-5H.

The patterns of Ca^+2^ release and MAPs recorded at −100 and −80 mV are similar to those recorded at −115 mM Na^+^_o_ (Figures 5E and 5F; compare with Figures 5A and 5B).

In fibers maintained at −60 mV, a slightly potentiated Ca^+2^ transient is seen in response to the first stimulus of the train; however, only every other later MAP recruited a Ca^+2^ transient afterwards, each of them reaching a peak similar to that of the Ca^+2^ transients recorded at −100 and −80 mV (Figure 5G, top panel). This pattern was seen in 2 out of 4 fibers. In contrast, skipping was never seen for MAP generation (Figure 5G, bottom panel).

As seen in 115 mM Na^+^_o_, when fibers are depolarized to −55 mV, a MAP is elicited by each pulse of the train. Nevertheless, in this case, MAPs are much smaller than those at 155 mM Na^+^_o_, peaking at about −10 mV (Figure 5H, top panel). As for experiments at 115 mM Na^+^_*o*_, only the first MAP triggers a Ca^+2^ transient at 40 mM Na^+^_o_ (Figure 5H, bottom panel).

#### 60 mM Na^+^_o_

In fibers exposed to 60 mM Na^+^_o_ (n=4), Ca^+2^ transients are recruited by each pulse of the trains as far as the membrane potential is held more negative than −80 mV (Figure 6A and 6B, top panels). The pattern of release along the train is similar to that seen at 115 and 90 mM Na^+^_o_, but no potentiation is seen in response to the first pulse at −80 mV. Likewise, in these conditions, robust MAPs are triggered at both membrane potentials, with OSs about 35 mV (Figure 6A and 6B, bottom panels).

**Figure 6.**
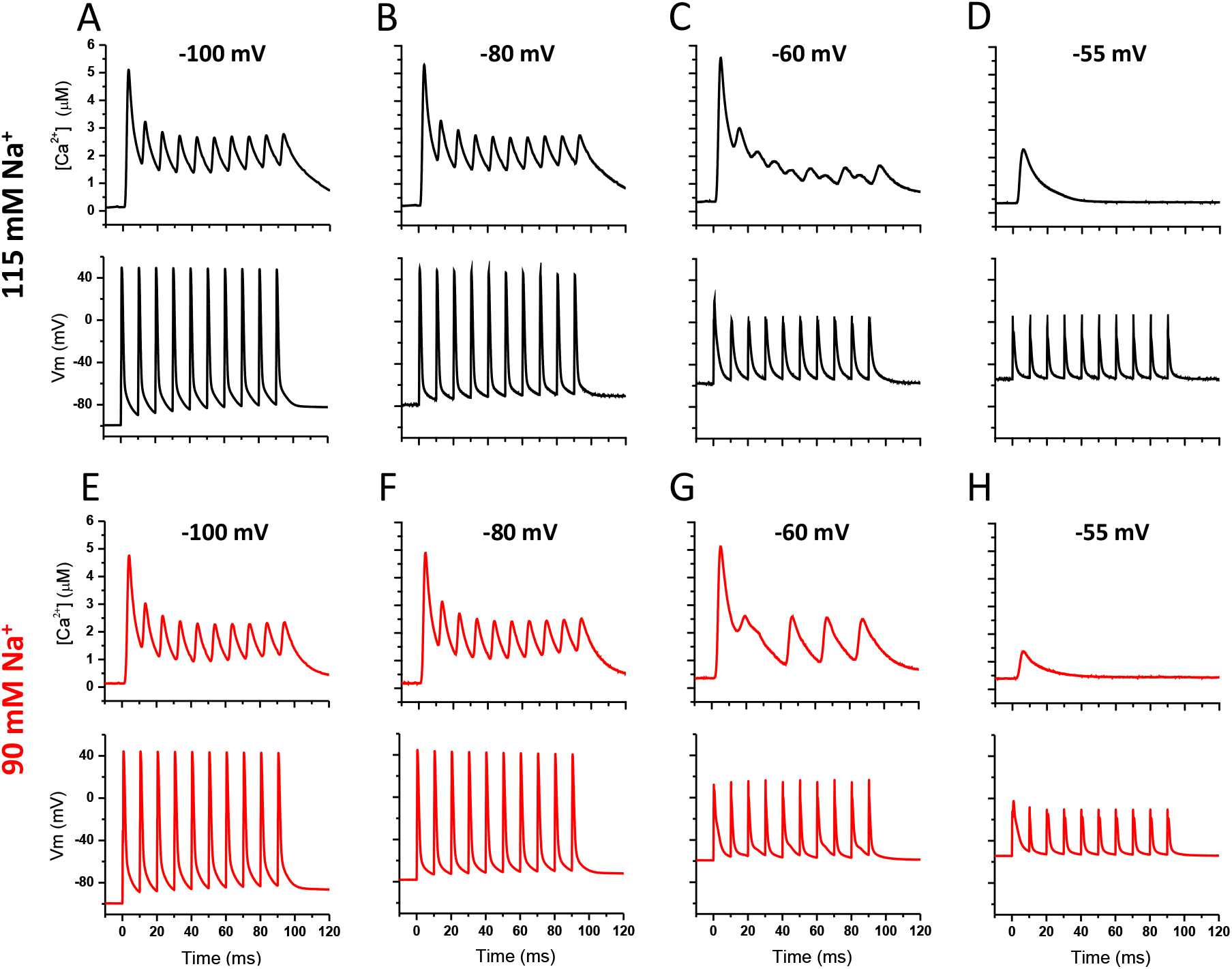
Effects of membrane potential on the Ca^+2^ release elicited by high frequency stimulation in fibers bathed in Ringer containing 115 or 90 mM Na^+^_o_. Top and bottom panels for each condition show the Ca^+2^ transients and their corresponding MAPs, respectively. Recordings were obtained from fibers maintained at the indicated holding potentials and exposed to either 155 (black traces) or 90 mM [Na^+^]_o_ (red traces). A 100 ms, 100 Hz train of supra-threshold pulses was used to stimulate the fibers. Similar responses were recorded in 4 different fibers for each 115 and 90 Na^+^_o_.

Nevertheless, at 60 mM Na^+^_o_ release along the train is not sustained at potentials as negative as −70 mV. In this case, the peak of each Ca^+2^ transient release decays along the train, and is barely noticeable as ripples by the 5^th^ or 6^th^ pulse (Figure 6C, top panel). In contrast, MAPs of preserved features, and OSs above 20 mV are generated in response of each pulse (Figure 6C, bottom panel). This results demonstrate that lowering Na^+^_o_ compromise the Ca^+2^ release but not the MAP generation along the trains. This is another example of an apparent disconnection between the amplitude of the MAPs and the recruitment of Ca^+2^ transients.

For further depolarizations to −65, small Ca^+2^ transients are only recruited in response to the first two pulses of the train, while robust MAPs are elicited by each stimulus (Figure 6D).

#### 40 mM Na^+^_o_

Reducing [Na^+^]_o_ to 40 mM almost completely inhibits Ca^+2^ release in fibers polarized between −100 and −70 mV. In most cases, after the second or third pulses, Ca^+2^ release is reduced to small ripples in response to each pulse of the trains (Figures 6E-6H, top panels). However, small Ca^+2^ transients are seen in response to the first pulse of the trains, their peaks had a similar voltage-dependence as that observed in response to single stimulation. Notably, at all the potentials explored, robust MAPs, with OSs of 20-30 mV, are elicited by each stimulus along the train (Figures 6E-6H, bottom panels). A similar depression of Ca^+2^ release was observed in other two fibers. As noticed above for single stimulus experiments, in our working conditions, extreme Na^+^_o_ deprivation decouples Ca^+2^ release from surface membrane depolarization (i.e. AMPs).

## Discussion

We have previously demonstrated the usefulness of our opto-electrophysiological method based on the use of inverted grease-gap chambers to study mechanisms of fatigue in segments of fibers from long muscles, which are not amenable for most electrophysiological techniques [7].

This approach allows for simultaneously studying excitability and Ca^2+^ homeostasis in rested fibers mechanically arrested by stretching (sarcomere length ~4 μm), while controlling the membrane resting potential and the intra- and extracellular milieus composition.

In our experimental conditions, non-propagating APs, i.e. MAPs, and Ca^2+^ transients free of mechanical artifacts can be recorded with minimal perturbations of the fiber’s Ca^2+^ buffering capacity. The use of a low affinity Ca^2+^ dye and in situ calibrations allow for quantitative fast time resolution studies of Ca^2+^ release [7].

Although muscle fatigue is multifactorial in origin, our capacity of studying the role of particular causative factors in isolation in rested fibers has proven to be mechanistically insightful [7].

In this work we assessed some of the tenets of the Na^+^ hypothesis for muscle fatigue. To this end we performed a study of the effects of steady state reductions in the [Na^+^]_o_ and depolarization on the electrical activity and Ca^2+^ release of fast skeletal muscle fibers exposed to (constant) physiological [K^+^]_o_, and expectedly, at constant [Na^+^]_i_ and [K^+^]_i_. Of relevance for data interpretation, it is expected that pre-stimulus membrane potential and [Na^+^]_o_ are the same at the surface and TTS membranes and that no radial membrane potential or [Na^+^]_o_ gradients exist. Consequently, pre-stimulus availability of Na^+^ channels and Na-EMF is the same at both membrane compartments.

Since most effects of raising [K^+^]_o_ on Ca^2+^ release are directly mediated by membrane depolarization [7], the combined effects of simultaneous alterations of [K^+^]_o_ and [Na^+^]_o_, typically used in force-measuring experiments, can be inferred from the interactions of the effects of Na^+^_o_ deprivation and membrane depolarization described here.

### Ca release in highly polarized fibers exposed to lowered [Na^+^]_o_

The main tenet of the Na^+^ hypothesis for muscle fatigue is that Na^+^_o_ depletion (and Na^+^i accumulation) will impair TTS AP propagation [11]. To what extent these concentrations should change to depress AP generation and propagation, and thus Ca^2+^ release, is unknown. Our experimental design may help defining what t-tubular [Na^+^], at constant [Na^+^]_i_ and in the absence of radial [Na^+^] gradients, may compromise the ECC process.

While interstitial [Na^+^] changes upon repetitive activation seems to be too small to support the Na^+^ hypothesis [13, 14]; and force in rested muscles is only reduced by very large [Na^+^]_o_ deprivation [11], these results do not fully rule out the Na^+^ hypothesis for muscle fatigue. Changes in [Na^+^] larger than those measured in plasma or interstitium are expected to occur in the TTS, particularly at high stimulation frequencies [5, 11, 15-17], and the change in [Na^+^]_o_ required to reduce force is smaller at high [K^+^]_o_ (i.e. at depolarized potentials) [11, 16, 18]. Moreover, simultaneous changes in [Na^+^]_i_ and [K^+^]_i_ will further reduce the Na-EMF and depolarize the fibers [3], as compared with the changes due only to Na^+^_o_ deprivation, as used in this work.

Although these studies provide insights on the causative relation between [Na^+^]_o_ deprivation and impaired force generation, the possible effects of [Na^+^]_o_ reductions on Ca^2+^ release has only been inferred so far.

Our single stimulus experiments at high membrane potential (i.e. −100 mV, Figure 1) demonstrated that Ca^2+^ release is practically immune to reductions of [Na^+^]_o_ to values as low as half the physiological concentration, but is significantly impaired when fibers are bathed in 40 mM Na^+^_o_ (Figure 1C). Our observations explain the effects of lowering [Na^+^]_o_ on twitch force of rested frog muscles as the twitch force-[Na^+^]_o_ relationship previously reported (Figure 2, [6]) and the peak Ca^+2^ transient-[Na^+^]_o_ relationship found here (Figure 3B) are very similar to each other.

Simultaneously recorded membrane potential shows that the grossly contrasting effects of extreme [Na^+^] deprivation on Ca^2+^ release does not concur with relatively minor changes in the OS (Figures 1 and 4).

There is then an obvious dissociation between the apparent intactness of MAPs and the pronounced impairment of the Ca^+2^ transients they triggered. In other words, at very low [Na^+^]_o_, the Ca^+2^ release mechanism seems to decouple from surface membrane APs (as reported by MAP recordings).

Our finding is not new; a similar dissociation, but between mechanical output and action potential amplitude was observed in frog fibers half a century ago by Bezanilla et al.; who concluded that “the action potential diminution alone is not sufficient to explain decreased tension” [15].

Our experimental paradigm is expected to assure that the surface and TTS membranes are enduring the same voltage and [Na^+^] transmembrane gradients (−100 mV and ~11.5Na^+^_o_:Na^+^_i_), which will not change in response to a single stimulation. Consequently, APs of similar features should be generated at both membrane compartments (i.e. similar to the magenta trace in Figure 1D); but if this was true, no significant Ca^+2^ release impairment was predicted at 40 mM Na^+^_o_. Since the recorded MAPs originate mainly from the surface membrane, our results suggests a limitation in the radial propagation of APs as the main mechanism underlying Ca^+2^ release impairment at high membrane potential and lowered [Na^+^]_o_, a condition affording a maximal availability of both Na^+^ channels and the voltage sensor for ECC.

A simple speculation is that at −100 mV, the t-tubule [Na^+^] should be reduced to 40 mM to impair TTS AP generation and conduction, a corollary is that, in physiological conditions, the same effects should be reached at a much lower [Na^+^]_o_ since [Na^+^]_i_ is expected to significantly increase during repetitive stimulation.

Indirect evidence from other laboratories also supports a possible failure of radial propagation of APs in reduced [Na^+^]_o_ [15, 18]. The larger diameter of frog fibers (~65 μm, this work) as compared with that of mouse fibers (40-50 μm, [19, 20]) should make the former more sensitive to low [Na^+^]_o_.

Simply stated, in its current implementation, our method clearly show that the Ca^2+^ release failure at 40 mM Na^+^_o_ and −100 mV cannot be explained on the basis of the observed MAP changes. Although our approach cannot provide direct insight on TTS AP propagation, instead, since ECC only occurs at the TTS membrane, the fact that impaired Ca^2+^ transients are recorded embodies in itself the evidence suggesting such defective radial propagation.

This challenging possibility deserves further investigations by means of potentiometric detection of TTS APs [21, 22]; and/or confocal detection of Ca^2+^ release [23, 24]; and recordings of Ca^2+^ release in electrically passive fibers under voltage-clamp conditions.

### Effects of membrane depolarization on Ca^+2^ release in fibers exposed to various [Na^+^]_o_

The experiments at different membrane potentials (Figures 2 and 3) confirmed our previous observations that depolarization has a dual effect on Ca^+2^ release triggered by single stimulation in fast frog muscle fibers exposed to physiological [Na^+^]_o_ [7]. In spite of ample differences in experimental approaches, a similar voltage dependence of Ca^+2^ release on membrane potential was also confirmed in murine skeletal muscles using different temperatures and Ca^+2^ sensors (30°C and Indo-1 [25]; 22°C and GCAMP6f [26]). The fact that intact fibers from long muscles (EDL and soleus) were used in these studies reinforces the validity of our approach using cut fibers.

Ca^+2^ release-Vm plots clearly shows a potentiation branch in the range of −100 to −60 mV; and a steep depression branch, at more depolarized potentials. The maximal potentiation occurs at the rather depolarized membrane potential of −65 mV (Figure 3, black trace and symbols). It is remarkable that Ca^+2^ release at −60 mV is comparable to that at −100 mV, and that robust Ca^+2^ transients, about 50% the amplitude of those at 100 mV, can still be recruited at −55 mV by MAPs having an OS of merely ~5 mV. Data from frog muscle fibers, obtained at credible space clamp conditions, show that Na channels’ availability is close to zero at −60 mV [27], and predicts that fibers should be mostly unexcitable at- and beyond this potential. Still, the possibility exist that the presence of CsF in the internal solution used in this work may have left shifted the voltage dependence of Na^+^ channels’ fast inactivation. Nevertheless, using normal [Ca^+2^]_o_ and Cs-aspartate instead of CsF in the internal solution, our simulations of Na^+^ currents for murine fibers suggest a similar voltage dependence of Na^+^ channels inactivation [28]. Thus, a discrepancy between data obtained in voltage- and current-clamp experiments exists, and needs to be resolved.

Here we have extended our previous study [7] to three more [Na^+^]_o_. Experiments at lower than physiological [Na^+^]_o_ led to three new findings.

A. A similar dual effect of depolarization on Ca^+2^ release was found at 90 mM Na^+^_o_, but in this condition, smaller potentiation at −65 mV and enhanced depression at −55 mV were seen. Clearly, those changes are due to Na^+^_o_ deprivation. Thus, reducing [Na^+^]_o_ somehow counteract the mechanism leading to Ca^+2^ release potentiation, and enhance the voltage-dependent mechanism that depress Ca^+2^ release.
B. We found that, at all voltages explored, the peak Ca^+2^ release is smaller in fibers exposed to 60 mM Na^+^_o_ as compared with that at 115 and 90 mM Na^+^_o_. Moreover, voltage-dependent potentiation, but not Ca^+2^ release depression, is lost at this [Na^+^]_o_ (Figure 2 and 3).
C. Interestingly, the depression branch of the voltage-dependence of Ca^+2^ release shifts leftward as [Na^+^]_o_ is lowered to 60 mM (Figures 3).

Altogether the results above demonstrate that Ca^+2^ release is more sensitive to depolarization the lower the [Na^+^]_o_.

Potentiation may be explained by the depolarization-dependent increase in resting [Ca^+2^]_i_ [7]. Our current observations are in agreement with previous data showing that conditioning subthreshold pulses and K-dependent depolarization can potentiate Ca^+2^ release and twitch tension [7, 25, 30, 31].

The lack of major effects of OS reduction on Ca^+2^ release may be, in part, explained on the basis of the voltage dependence of Ca^+2^ release (as measured in voltage clamp conditions), which saturates at about 20 mV [9, 10, 20]. Thus, in principle reducing the OS from 50 mV to ~20 mV may cause minor changes on the amplitude of Ca^+2^ transients. Therefore, the difference between the Vm at which the Ca^+2^ release saturates and the OS represents a large safety factor for AP triggered ECC.

Several mechanisms can lead to Ca^+2^ release depression, such as MAP impairing, radial propagation failure or inactivation of the ECC mechanism itself. Na^+^ channel inactivation and reduced Na-EMF may underlie changes in MAPs and radial propagation. Reduced OS at 60 mM Na^+^_o_ (Figure 4) may overcome the effects of raised [Ca^+2^]_i_, thus preventing potentiation.

Drastic reduction of [Na^+^]_o_ to 40 mM dwarfs Ca^+2^ release at all membrane potentials tested (Figure 3). Remarkably, in this condition, relatively large MAPs are recorded between −100 and − 70 mV (Figure 4), suggesting a radial conduction failure as the cause of Ca^+2^ release impairment. These findings at 40 mM Na^+^_o_ extend those discussed above and uncover an apparent decoupling of Ca^+2^ release from MAPs at all membrane potentials tested.

For each [Na^+^]_o_ tested, depolarization from −100 to −70 mV have modest effects on Ca-FWHM, but in this range of membrane potentials, Ca^+2^ transients at 60 and 40 mM Na^+^_o_ are longer lasting than those at 115 and 90 mM Na^+^_o_. The Ca-FWHM at these two later [Na^+^]_o_ were similar. The increased Ca-FWHM found at membrane potentials more positive than −70 mM (Figure 3) may help sustaining contractions in depolarized fibers.

### Effects of lowering [Na^+^]_o_ on Ca^+2^ release in fibers polarized at various membrane potentials

The dependence Ca^+2^ release on [Na^+^]_o_ is summarized in Figure 3B. Reductions of [Na^+^]_o_ to 90 mM (~20% below the physiological value) have no significant detrimental effects on Ca^+2^ release as far as the fibers are maintained at membrane potentials more negative than −60 mV (Figure 3B, all traces). In fact, as seen in normal Ringer, potentiation of Ca^+2^ release ensues at this [Na^+^]_o_ along this voltage range. This demonstrates that, in the scenario of single stimulation, the ECC process has a large safety factor for both membrane depolarization (a 40 mV change) and Na^+^ depletion (a 25 mM change), regardless these changes occur in isolation or in combination. In contrast, a larger reduction in Ca^+2^ release is found in response to a further 5 mV depolarization in fibers bathed in 90 mM Na^+^_o_ as compared with that found at 115 mM Na^+^_o_. This results shows that Na^+^_o_ deprivation worsens the large depression of Ca^+2^ release seen at 115 mM Na^+^_o_ and −55 mV. This additional effect is due to the lower [Na^+^]_o_ and is independent of the availability of Na^+^ channels and ECC voltage sensor at −55 mV. Comparable [Na^+^]_o_ changes are not expected to occur in either the plasma or the interstitial spaces of muscles stimulated repetitively [11], nonetheless, similar reductions in TTS luminal [Na^+^] has been predicted to occur in <0.5 s in frog muscle fibers stimulated at 60 Hz (Figure 6 in [15]).

Even halving the physiological [Na^+^]_o_ results in only a small reduction of Ca^+2^ release in fibers polarized to potentials more negative than −70 mV (Figure 3B); demonstrating that the ECC process operates within a large voltage safety factor when recruited by single stimulation in rested fibers. Nonetheless, depolarizations to −65 and −60 mV greatly impair ECC (Figure 3B, cyan and magenta traces). This result clearly shows that furthering [Na^+^]_o_ deprivation increases the sensitivity of the ECC process to depolarization. The detrimental effects of Na^+^_o_ deprivation is exemplified by the fact that membrane potentials affording potentiation at 115 and 90 mM Na^+^_o_, lead to significant depression at 60 mM Na^+^_o_.

At extreme low values (i.e. 40 mM), [Na^+^]_o_ becomes the limiting factor for ECC activation as shown by the abrupt depression of the Ca^+2^ release elicited at all potentials (Figure 3B, all traces). This manifests as an apparent decoupling of Ca^+2^ release from robust surface MAPs elicited from −100 to −65 mV.

Overall, our data demonstrate that in fast frog fibers ECC operates safely within large margins of membrane potentials and [Na^+^]_o_ changes; and that independent or combined reductions of both parameters beyond safety limits impair Ca^+2^ release. In this way, depolarization renders the ECC process more sensitive to lowered [Na^+^]_o_; and conversely, at lowered [Na^+^]_o_, Ca^+2^ release depression occurs in response to smaller depolarizations.

The combined effects of both perturbations on Ca^+2^ release are complex (see Figure 7) and seem to be more than additive. Synergistic effects of increasing [K^+^]_o_ (akin to current injection driven depolarization) and reducing [Na^+^]_o_ on the mechanical output of rested muscles were first reported for frog muscles [11]. Here we present the first demonstration of similar effects of depolarization (mimicking the effects of raised [K^+^]_o_ [7]) and [Na^+^]_o_ deprivation on Ca^+2^ release. Our data show that the synergistic effects of [K^+^]_o_ augmentation and [Na^+^]_o_ deprivation on twitch force are mediated by similar effects on Ca^+2^ release exerted by depolarization and [Na^+^]_o_ deprivation. It should be noticed, nevertheless, that K^+^_o_ exerts direct effects on ion transport mechanisms, relevant to AP conduction in the TTS, which are not mimicked by depolarization (this work). Raised K^+^_o_ increases the conductance of the potassium inward rectifier channels [32, 33]. While this has the beneficial effect of increasing K^+^ reuptake from the TTS [32], it also have the detrimental effect of reducing the space constant of the TTS, further compromising radial propagation. Increased K^+^_o_ upregulated NaK-pump activity [34, 35]. This may help limiting changes in transmembrane K^+^ and Na^+^ gradients during activity [36], thus limiting fatigue development.

**Figure 7.**
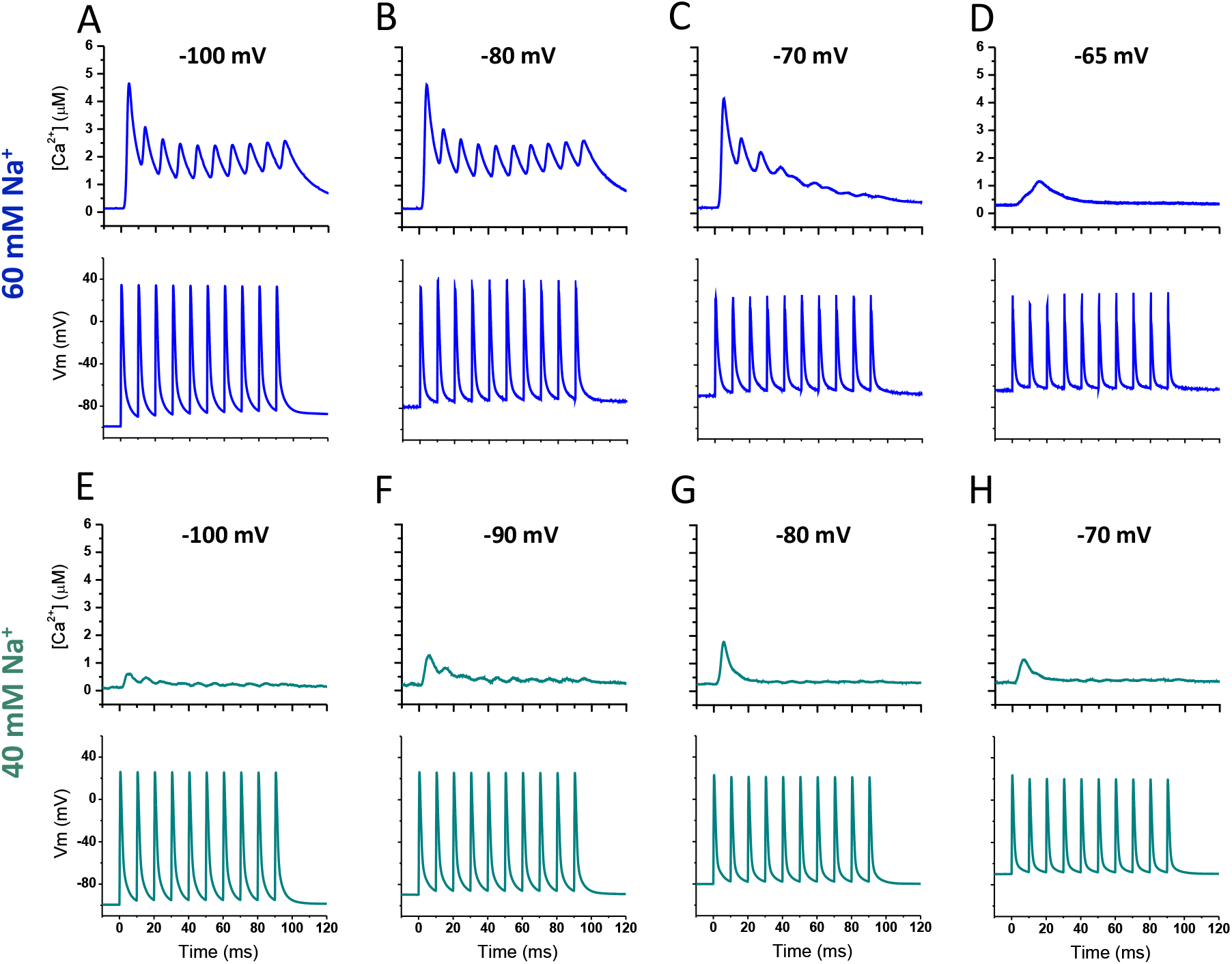
Effects of membrane potential on the Ca^+2^ release elicited by high frequency stimulation in fibers bathed in Ringer containing 60 or 40 mM Na^+^_o_. Top and bottom panels for each condition show the Ca^+2^ transients and their corresponding MAPs, respectively. Recordings were obtained from fibers maintained at the indicated holding potentials and exposed to either 60 (blue traces) or 40 mM [Na^+^]_o_ (green traces). A 100 ms, 100 Hz train of supra-threshold pulses was used to stimulate the fibers. Similar responses were recorded in 4 different fibers for 60 and 3 fibers for 40 Na^+^_o_.

### The dependence of Ca^+2^ release on [Na^+^]_o_ and Vm explains the effects of changes in [Na^+^]_o_ and [K^+^]_o_ on twitch force

The resemblance between the dependence of Ca^+2^ release on resting membrane potential determined at various [Na^+^]_o_ (Figure 3A, this work) and the dependence of twitch force on [K^+^]_o_ and [Na^+^]_o_ (Figure 2A in [11]) is quite remarkable.

For single ion changes it was found [11] that at normal [K^+^]_o_, twitch force is practically insensitive to changes in [Na^+^]_o_; while at normal [Na^+^]_o_, small increases in [K^+^]_o_ potentiates twitch force, but large increases abolish force generation. Lowering [Na^+^]_o_ reduces the potentiation effect of raising [K^+^]_o_. Moreover a synergistic interaction between the effects of changes in both ions was found. Lowering [Na^+^]_o_ reduces the force potentiation effect of increasing [K^+^]_o_ and reduces the [K^+^]_o_ needed to depress force; i.e. the depression branch of the force-[K^+^]_o_ relationship shifts to the left. Finally, almost no force potentiation is seen at intermediate [Na^+^]_o_.

We found analogous changes in Ca^+2^ release in response to equivalent alterations of membrane potential and [Na^+^]_o_, demonstrating that twitch force potentiation or depression are mediated by equivalent changes in Ca^+2^ release.

Similarly, at physiological membrane potentials, the Ca^+2^ release (Figure 3B, this work) and the peak twitch tension show the same dependence on [Na^+^]_o_ (Figure 2A in [6]). This demonstrates that in both frog and mouse the attenuation of peak twitch force observed at very low [Na^+^]_o_ is mediated by a depression of Ca^+2^ release.

Overall, the changes in twitch force in response to individual or combined alterations of [Na^+^]_o_ and [K^+^]_o_ are closely mirrored by changes in Ca^+2^ release in response to Na^+^_o_ deprivation (at constant Vm) and membrane depolarization (at constant Na^+^_o_). Our current data suggests that the combined effects of raised [K^+^]_o_ and reduced [Na^+^]_o_ on twitch force are independent, at least in part, from the presence of K^+^ per se, but are instead mediated by K-dependent membrane depolarization [7].

Consequently, our study is the first direct demonstration that the modulation of twitch force generation by changes in [Na^+^]_o_ and/or [K^+^]_o_ is mediated by the effects that those changes exert on Ca^+2^ release. Before our work, the mechanism underlying force dependence on Na^+^_o_ and K^+^_o_ could only be anticipated [11]. The next question to answer is how depolarization (or raised K^+^_o_) and/or Na^+^_o_ deprivation limits Ca^+2^ release?

### Voltage- and [Na^+^]_o_-dependent MAP overshoot changes do not correlate with changes in Ca^+2^ release

It is well stablished that the ECC process in skeletal fibers is only recruited, in a graded and saturable fashion, by TTS membrane depolarization. Since in physiological conditions Ca^+2^ release is triggered by Na^+^-dependent APs propagating radially in the TTS, it is conceivable that factors altering the generation or waveform (e.g. amplitude, OS, duration) of those APs will in turn alter Ca^+2^ release. Consequently, understanding how K^+^-dependent depolarization and Na^+^_o_ deprivation affect the generation and waveform of APs may help explaining how K^+^_o_ accumulation (membrane depolarization) and Na^+^_o_ deprivation affects Ca^+2^ release and ultimately force generation. We first looked at the voltage- and [Na^+^]_o_-dependence of the OS and compared them with those of the Ca^+2^ release.

Our previous (Figure 5C in [7]) and current data (Figure 4) concur with previously published relationships between AP OS and [K^+^]_o_ (Figure 5 in [4]), [Na^+^]_o_ (Figure 6 in [6]), and membrane potential (Figure 2E in [26]). Here we have extended those studies to a larger number of conditions.

According to the Na^+^ hypothesis for muscle fatigue [6] impaired mechanical output can be explained on the basis of a reduction of Na-EMF predicted from Na^+^_o_ depletion (mainly at the TTS lumen) and Na^+^_i_ accumulation during sustained activity. Since the ENa sets the maximal depolarization reachable during an AP, this will translate to a reduced OS. At constant [Na^+^]_o_, the OS is also reduced by depolarizations through voltage-dependent inactivation of Na^+^ channels.

Accordingly, independently or in combination, depolarization in our case (or increased [K^+^]_o_ in other studies) and Na^+^_o_ deprivation are expected to reduce Ca^+2^ release via a reduction of the OS. Our results from single stimulation experiments does not concur with this prediction, instead a complex relationship between Ca^+2^ release and OS was determined (Figure 8, see also Figure 9). The insensitivity of Ca^+2^ release to OS changes at various [Na^+^]_o_ and membrane potentials suggests it is not the only factor determining the Ca^+2^ transient peak. Others have previously shown that force generation is relatively insensitive to reduced OS resulting from partial Na^+^ channel blockage or reduced ENa at low [Na^+^]_o_ [6, 15, 37].

**Figure 8.**
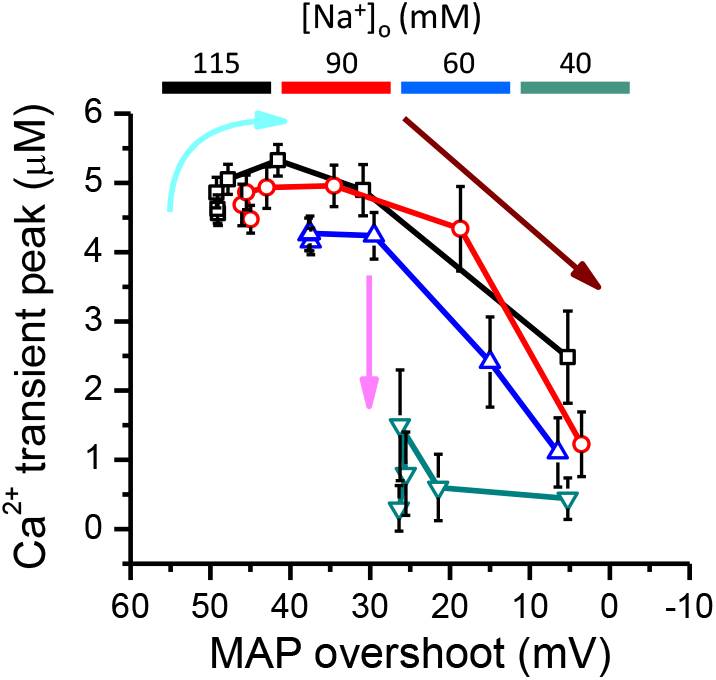
Relationship between the peak of Ca^+2^ transients and the overshoot of MAPs. The peak of Ca^+2^ transients are plotted as a function of the OS. Lines and symbols represent data obtained at 115 (black line and symbols), 90 (red line and symbols), 60 (blue line and symbols) and 40 mM Na^+^_o_ (green line and symbols). The cyan arrow indicates potentiation of Ca^+2^ transients as the OS is reduced from 50 to 40 mV. The magenta arrow indicates the depression of Ca^+2^ transients at 40 mM Na^+^_o_ and large OSs. The brown arrow indicates a monotonic relationship between the Ca^+2^ transient’s peak and MAP FWHM.

**Figure 9.**
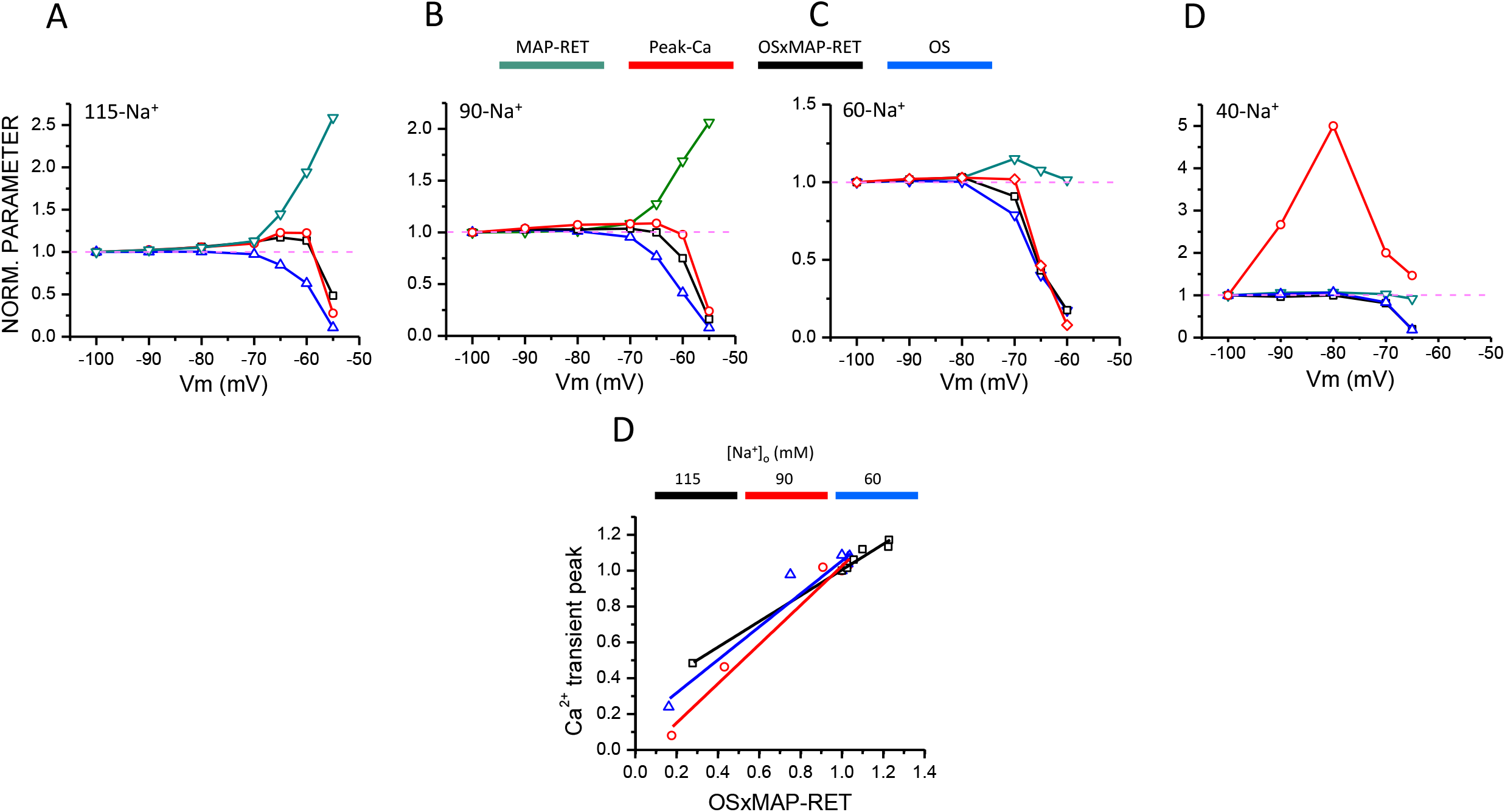
Correlation between the peak of Ca^+2^ transients and OSxMAP-RET. **A-D.** The MAP-RET, the peak of the Ca^+2^ transients (Peak-Ca^+^), the OS and the product of the OS times MAp-RET (OSxMAP-RET) measured at 115, 90, 60 and 40 mM Na^+^_o_ were normalized and plotted as a function of the holding potential (Vm). The [Na^+^]_o_ is color coded as indicated by the colored bars. Each parameter was normalized to its value at −100 mV (indicated in each panel by the dashed lines). **E.** Correlation between the peak of Ca^+2^ transients and OSxMAP-RET. The peak of Ca^+2^ transients determined at 115, 90 and 60 mM [Na^+^]_o_ are plotted as a function of the corresponding OSxMAP-RET. The data points for each [Na^+^]_o_ were fitted to linear regressions. The R^2^ values were 0.99, 0.94, 0.98 and 0.47 for data obtained at 115, 90, 60 and 40 mM [Na^+^]_o_, respectively. The data for 40 mM [Na^+^]_o_ are not shown. The [Na^+^]_o_ is color coded as indicated by the colored horizontal bars.

In our case, two extreme experimental conditions exemplify the apparent anomalous relationship between the OS and Ca^+2^ release elicited by single stimulation (Figure 8); a) Ca^+2^ release is potentiated in conditions that reduce the OS (e.g. −75 to −60 mV, 115 to 90 mM Na^+^_o_; Figure 8, cyan arrow). In contrast, the OS depends monotonically on both depolarization and lowered Na^+^_o_ (Figure 4); b) Ca release is almost abolished in conditions that allow for large OSs (e.g. −100 to − 70 mV, 40 mM Na^+^_o_; Figure 8, magenta arrow). In other conditions (−70 to −55 mV, 115 to 60 mM Na^+^_o_; Figure 8, brown arrow), the peak of the Ca^+2^ transients declines monotonically as OS is reduced.

The observed dissociation between OS and Ca^+2^ transient peak may be wrongly interpreted as a decoupling between TTS APs and Ca^+2^ release, but this is only apparent; a distinction needs to be made between MAPs (or longitudinally propagated APs) and TTS APs as this fact is usually overlooked. Many published evidences suggest that OS is not an appropriate figure of merit to predict Ca^+2^ release or contraction [6, 7, 15, 37]. Since MAPs and ECC ocurrs at different membranes, it is possible that MAPs or longitudinally propagated APs can be elicited in conditions limiting or preventing AP propagation along the TTS membranes, as seen, for example, at 40 mM Na^+^_o_ (or repetitive stimulation, see below). In this conditions, robust MAPs can be elicited but only small Ca^+2^ transients are recruited. While impaired Ca^+2^ release attests in itself to a TTS AP propagation failure, a direct test of this possibility is missing. An alternative condition, preventing longitudinally propagated APs or MAPs but allowing radial propagation is difficult to conceive. In physiological conditions, longitudinal propagation is the only way to convey APs initially generated at the neuromuscular junction to t-tubule openings.

Intrinsic differences between the surface and TTS membranes may be at the root of the seemingly differential sensitivity of both compartments to the same depolarization and Na^+^_o_ deprivation, thus explaining TTS AP propagation failure in the presence of robust MAPs. The TTS is a diffusion-limited convoluted compartment with mostly unknown space constant and luminal conductivity. The TTS is delimited by a membrane of different lipid composition and lipid order as compared with those of the surface membrane [38, 39]; and lipids (e.g. cholesterol) are known to modulate ion channel activity [40]. The actual endowment of ion transport systems in the TTS is still a matter of debate (see for example [41-43]), but most probably, it is different from that of the surface membrane in terms of channel density, stoichiometry, and isoforms present. For example, different isoforms of the NaK-pump are present at each membrane compartment of murine muscle fibers, making them physiologically different [34, 44]. Although only one Na^+^ channel isoform prevails in adult fibers (NaV_1.1_), Na^+^ channels at the TTS and surface membranes may work differently due to differences in lipid environment.

It should be noted that, at the level of single fibers, radial gradients of ion concentrations and membrane potential are not expected to occur in our single stimulation experiments, but may play a key role during sustained activity. In physiological conditions, radial gradients of membrane potential and ion concentrations predict close to normal APs at the surface membrane but impaired TTS APs, worsening towards the fiber center.

One important factor for the generation and conduction of APs, particularly in the TTS, usually overlooked is the “resting” (or pre-stimulus) membrane conductance [45]. The main ionic conductances contributing to the resting conductance are expected to increase during sustained activation. The chloride conductance will increase due to depolarization per se [42], and the potassium inward rectifier conductance will increase due to its dependence on [K^+^] [32]. Both of this factors will reduce the space constant of the TTS, compromising the generation and conduction of APs.

### MAP duration and Ca^+2^ release

The MAP duration is also expected to affect the features of Ca^+2^ release. We used two parameters to assess the MAP duration, the FWHD and the RET (see Methods). The first is a standard way of measuring the duration of waveforms, the second was conceived from the classical mechanical effective time [11, 12] and the membrane potential threshold for Ca^+2^ release [9, 10].

The MAP-FWHM measured here (Figure 5A) showed a voltage dependence similar to that previously reported (Figure 5D in [7]). Nevertheless, while we found that for all [Na^+^]_o_ used, MAPs shorten as fibers are depolarized from −100 to −70 mV, it was previously reported that AP FWHM is almost insensitive to depolarization along a (predictable) similar range of potential (Figure 5B in [4]).

Since, unexpectedly, the voltage dependence of MAP-FWHM and Ca^+2^ release seem to be negatively correlated (compare Figures 3A and 5A), we resorted to measure MAP-RET (Figure 5B). While in the range of −100 to −70 mV, MAP RET better correlate with Ca^+2^ release, it still show a negative correlation with Ca^+2^ release for further depolarization (Figure 5B).

Since the OS and the RET have similar apparent voltage independence between −100 and −70 mV, but opposite voltage dependence in the range of −70 to −55 mV, we reasoned that the product of both parameters (OSxMAP-RET) may better correlate with Ca^+2^ release (Figure 9). In fact, we found that the voltage dependence of OSxMAP-RET closely approximate the voltage dependence of Ca^+2^ release at [Na^+^]_o_ between 115 and 60 mM (Figures 9A-9C). As expected no correlation was found at 40 mM Na^+^_o_. This was confirmed by plotting the Ca^+2^ transient peaks as a function of OSxMAP-RET determined at the same Vm and [Na^+^]_o_. A high correlation was determined for the data at 115-60 mM Na^+^_o_ (Figure 7E, see legend for R^2^ values), suggesting a causative relationship between both parameters. A similar correlation between the peak of Ca^+2^-dependent fluorescence transients and the area of APs was recently described [26].

### Na^+^_o_ deprivation and depolarization impair Ca^2+^ release in rested fibers stimulated at 100Hz

Tetanic force is more sensitive to [Na^+^]_o_ deprivation or increased [K^+^]_o_ than twitch tension, and this two factors have been shown to synergistically impair tetanic force generation [11]. A similar study of Ca^2+^ release in these conditions was missing, thus we measured, in a relatively small number of fibers, Ca^2+^ transients elicited by short, 100 Hz trains in rested fibers exposed to 115-40 mM Na^+^_o_ and maintained at holding potentials between −100 and −55 mV (Figures 6 and 7). Our approach is able to resolve the Ca^2+^ release events elicited by each pulse of the trains, providing a continuous quantitative readout of [Ca^2+^]_i_ changes and simultaneous recordings of the underlying MAPs.

Notably, our data demonstrate that fibers chronically exposed to [Na^+^]_o_ as low as half the physiological value can generate close to normal Ca^2+^ transients at a frequency of 100 Hz as far as the membrane potential is more negative than −80 mV.

In this conditions, robust MAPs are elicited and, in agreement with single pulse experiments, potentiation of the first Ca^+2^ transient of each train is seen at 115 and 90 mM Na^+^_o_, but not at 60 mM Na^+^_o_ (Figure 6-7).

At this range of membrane potentials a high gNa is expected, consequently the relatively small reduction of the OS observed is mainly due to the imposed reduction in ENa. The results suggest that gNa is the predominant factor governing the ECC process in this conditions, for this reason Ca^+2^ release seems immune to reduced [Na^+^]_o_.

Interestingly, at 115 and 90 mM [Na^+^]_o_, Ca^+2^ release from the second pulse is compromised, regardless the first Ca^+2^ transient is potentiated, if the fibers are depolarized to −60 mV (Figures 6C and 6G). At this potential a reduced gNa is expected; and this may explain why MAPs have smaller OSs. However, this cannot explain why the pattern of Ca^+2^ release is irregular while that of MAP generation is not. The results suggests that, notwithstanding [Na^+^]_o_ is high, at potentials that predicts a reduced gNa, the ECC process cannot respond normally at high frequency. The abnormal response to high frequency stimulation and the fact that the first Ca^+2^ transient of the train is normal is another example of an apparent disconnection between Ca^+2^ release and MAP features. One possible explanation is that the surface and TTS membranes respond differentially to the repetitive stimulation. In practical terms, these findings suggest that during repetitive stimulation the propagation mechanism along the TTS or the Ca^+2^ release mechanism has a lower safety factor in comparison to that observed in response to single stimulation.

Lowering [Na^+^]_o_ to 60 mM further compromise the response of the ECC process to tetanic stimulation, since Ca^+2^ release failure is observed at more negative membrane potentials (i.e. −70 mV), no matter the MAPs has a much larger amplitude than that of those recorded at −65 mV in fibers exposed to 115 and 90 mM Na^+^_o_ (Figure 6C and 6D). A larger gNa, as expected at −70 mV, may explain why the MAPs are larger at a lower [Na^+^]_o_, but is at odds with the ECC failure. Therefore, lowering [Na^+^]_o_ to 60 mM seems to further reduce the safety factor for Ca^+2^ release during repetitive stimulation. In these conditions, it seems that the [Na^+^]_o_ is the predominant factor determining the apparent decoupling of the ECC process from (surface membrane) MAPs. As expected from single pulse stimulation, Ca^+2^ release failed at all potentials when extremely low [Na^+^]_o_ even though MAP have large OSs (Figure 6E-6H).

As for single stimulation, we find that the OS per se fail to predict Ca^+2^ release in most conditions explored. The most dramatic demonstration of this is seen at 40 mM Na^+^_o_, robust MAPs, with OSs between 20 and 30 mV, failed to elicit normal Ca^+2^ release even in response of the first pulse of the trains. For [Na^+^]_o_ from 115 to 60 mM, Ca release failure at extreme depolarization was not due to failing MAP generation (Figures 5D and 6H, Figure 7D). Ca release can either be defective or fail along the trains while non-skipping, regenerative responses (MAPs) may peak between −5 and 20 mV are simultaneously recorded.

It should be noted that, in agreement with force measurements [11], aside for the response to the first stimulus, at none of the conditions studied, Ca^+2^ release was potentiated during the trains; all the experimental conditions tested resulted in no effect or depression of Ca^+2^ release.

Overall, our results demonstrate that Ca^+2^ release in response to repetitive stimulation is more sensitive to depolarization (and thus, to increased K^+^_o_) and Na^+^_o_ deprivation than Ca^+2^ release elicited by single stimulus. Moreover, as for single stimulation, when imposed in combination, depolarization and Na^+^_o_ deprivation seem to synergistically impair Ca^+2^ release in response to repetitive stimulation; as reported for tetanic force force [11]. A likely explanation for the effects of these two factors is a depression of the generation and conduction of APs along the TTS. However, possible alterations of other steps of the ECC during repetitive activation needs to be determined.

Our data provide, for the first time, a mechanism to explain how Na^+^_o_ deprivation and K^+^_o_ accumulation cause depression of tetanic force, and why K^+^_o_ accumulation potentiates twitch force but does not cause tetanic force potentiation.

### Further directions

The present work extends our previous study on the influence of changes in the [K^+^]_o_ and membrane depolarization on Ca^+2^ release [7]. Although amphibian muscles were used, both works provided novel insights on the mechanisms of muscle fatigue in particular and ECC in general. We plan to perform a similar study in myosin-typified mammalian muscle fibers maintained at close to physiological conditions. Combining electrophysiological and optical techniques [21, 32] and working in both current- and voltage-clamp conditions, the immediate goal is to directly assess the generation and propagation of APs along the TTS and the response of subsequent events of the ECC process to repetitive stimulation and changes in external ion concentrations and membrane potential. The long term goal is to accomplish similar studies in human muscle fibers. We have already demonstrated the feasibility of using our experimental approach to study the ECC process in fibers obtained from human biopsies [46], which are otherwise inaccessible to most standard electrophysiological techniques.

### Summary

The effects of depolarization and Na^+^ deprivation of Ca release elicited by single stimulation can be summarized as follows: a)Depolarization has a dual effect on Ca^2+^ release. At physiological [Na^+^]_o_, depolarizations to values more negative than −60 mV produce Ca^2+^ release potentiation, and depolarizations to values more positive than −60 produce a steep depression of Ca release. b)Na^+^_o_ deprivation has a monotonic detrimental effect on Ca^2+^ release. Potentiation is reduced at 90 mM Na^+^_o_ and eliminated at 60 mM Na^+^_o_. At both [Na^+^]_o_, voltage-dependent depression of Ca^2+^ release occurs at membrane potentials more negative than −60 mV. At 40 mM Na^+^_o_, Ca^2+^ release is partially decoupled from MAPs.

At [Na^+^]_o_ larger than 60 mM fibers can produce normal Ca^2+^ transients at 100 Hz if polarized to membrane potentials more negative than −80 mV. Larger depolarization impair Ca release along 100 Hz trains. In fibers polarized between −100 and −70 mV, Ca^2+^ release decouples from close to normal trains of MAPs, when exposed to 40 mM Na^+^_o_.

The effects of depolarization and Na^+^_o_ deprivation synergistically depress Ca^2+^ release.

The OS and duration of MAPs are not figures of merit to predict Ca^2+^ release; the parameter OSxMAP-RET better correlates with the peak of Ca^2+^ transients.

We proposed that a compromised TTS AP generation and/or conduction, not MAP impairment, may explain the detrimental effects of depolarization and Na^+^_o_ deprivation on Ca^2+^ release.

We conclude that the effects of increased K^+^_o_ and Na^+^_o_ deprivation on twitch and tetanic force generation of rested fibers can be explained on the basis of the effects of membrane depolarization and Na^+^_o_ deprivation on Ca^2+^ release.

## Acknowledgments

This work is part of MQ Ph.D dissertation Thesis.

